# Viral capsid delivery of cGAMP enhances STING-dependent antitumor immune response

**DOI:** 10.64898/2026.06.26.734859

**Authors:** Paul Huang, Yeara Jo, Hannah S. Martin, Rutger D. Luteijn, David H. Raulet, Matthew B. Francis

## Abstract

Therapies to activate the STING immune response pathway represent promising potential anticancer treatments. However, the native STING activating molecule, 2’,3’-cGAMP, is a poor drug candidate due to its susceptibility to nuclease degradation and its relatively poor cell uptake. In this study, we present a nanoscale delivery vehicle based on the bacteriophage MS2 virus-like particle that can both protect cGAMP and deliver it into cells to access and bind cytosolic STING. MS2-delivered cGAMP achieved greatly increased STING activation potency relative to both free cGAMP and a nuclease-resistant synthetic cGAMP analog. In an *in vivo* murine colon carcinoma model, MS2-cGAMP elicited significant and prolonged antitumor activity in a STING-dependent manner at 50-fold lower concentrations relative to free cGAMP and synthetic analogs. These results demonstrate that MS2 delivery of cGAMP can yield a highly potent STING agonist immunotherapy with *in vivo* anticancer activity.

## INTRODUCTION

The generation of effective antitumor immune responses is key to the treatment of highly aggressive cancers, especially in those that have shown limited benefit from conventional chemotherapies^1^. This can be done through stimulation of relevant innate immune pathways in the tumor microenvironment that signal the recruitment and activation of adaptive immune cells. One such pathway is the stimulator of interferon genes (STING) pathway, whereby damaged or pathogen-infected cells sense cytosolic DNA and initiate an innate immune response, secreting type I interferon and other proinflammatory signals into the extracellular milieu^2–4^. These in turn can both activate NK cell killing activity and act on T cells and antigen-presenting cells to elicit CD8 and CD4 T cell responses^3–5^. As a result, activation of the STING pathway is an important early link between the innate and adaptive immune systems.

STING pathway activation has been implicated in various antitumor responses and long-lasting antitumor immunological memory^6–8^, which has led to much work in developing anticancer therapeutics to activate STING^9–12^, albeit with limited clinical success to date. Many of these STING agonist drug candidates mimic the structure and binding affinity of the native STING agonist, cyclic GAMP (cGAMP), which is produced by the enzyme cyclic GMP-AMP synthase (cGAS)^13,14^. However, while cGAMP is a small molecule that binds and activates STING with nanomolar affinity, it exhibits poor pharmacological properties as a free drug due to its negatively charged phosphate groups, the bonds of which are susceptible to degradation by phosphatases^15^ and which impart poor cell membrane permeability. Indeed, regulation of proteins that control cGAMP concentration and cellular localization, such as extracellular phosphatases and cell-surface anion transporters, is an important mechanism through which STING activation is naturally modulated in the body^16,17^. Dysregulation of these mechanisms in cancers can in turn suppress STING activity^18^.

To counteract these challenges, initial drug design efforts for STING agonists focused on conferring more optimal “drug-like” properties to cyclic dinucleotides (CDNs) with high structural similarity to native cGAMP. These employed phosphorothioate analogs of CDNs to be more resistant to known cGAMP-degrading phosphatases such as ENPP1^15^. However, these structures still retain their anionic character, making them poorly membrane permeable and reliant on the same heterogeneously expressed cell-surface transporters as native cGAMP^19,20^; furthermore, phosphorothioate drugs have clearance pathways of their own^21^. The resulting low potency and poor cellular uptake of CDN drugs requires their administration at relatively high dosages to achieve therapeutic responses, increasing the likelihood of unfavorable side effects from off-target responses^22^. With these inherently poor pharmacokinetics, cGAMP mimics have fared poorly in clinical trials^9,23^. With increased structural understanding of how cGAMP binds and activates STING, later STING agonists have used more diverse, “drug-like” functional groups, such as a class of diaminobenzimidazoles (diABZI), to achieve low-nanomolar STING activation^24–27^. However, many non-nucleotide STING agonists have more modest binding affinities, and the toxicity profiles and off-target effects of these structures are less predictable.

Given the difficulty in designing a STING agonist drug with high bioavailability, potency, and specificity, some efforts have turned to nanoscale drug delivery platforms to optimize the administration of native cGAMP to cells of interest. In many cases, the anionic nature of cGAMP has been leveraged by using electrostatic interactions to associate the drug to delivery vehicles with cationic nanomaterials, such as polymer-based micelles, vesicles, and other nanoparticles, as well as membrane-enveloped virus-like particles^28–35^. While these structures protect cGAMP and facilitate cell entry, the non-covalent association of cGAMP to the nanoparticles complicates drug loading consistency and leaves the potential for leaky release of the drug. While this can be solved by covalent attachment, most linkage chemistries require a modified CDN structure^36–38^, which can affect STING binding affinity. Additionally, many nanoparticles do not efficiently localize their drug cargo to the cell cytosol, where STING is located, resulting in only modest increases in STING activation efficacy. Therefore, we sought to develop a delivery platform that could covalently attach native cGAMP and release it tracelessly inside of cells with high efficiency.

We previously reported that a virus-like particle (VLP) based on bacteriophage MS2 bearing two cationic amino acid mutations exhibited enhanced cellular uptake^39^ and delivery of anionic drug cargo, which we used to deliver a phosphorothioate CDN STING agonist to cells^40^. The CDN was attached to the porous interior of the capsid by way of a reductively cleavable disulfide linker. Here, we modified native cGAMP analogously with a self-immolative linker, enabling covalent attachment in the viral capsid interior without compromising the native binding affinities of cGAMP. We further showed that MS2 VLP delivery of cGAMP yields an enhanced STING response relative to free CDNs, a result of efficient delivery into the cell as well as protection of cGAMP from extracellular phosphatases. This potency enhancement translated to an *in vivo* murine carcinoma model, whereby MS2-delivered cGAMP was demonstrated to elicit a STING-dependent antitumor response at a 50-fold lower dosage level compared to free cGAMP or a phosphorothioate CDN drug. These results demonstrate a highly effective drug delivery platform for the administration of cGAMP as a potent anticancer immunotherapy.

## RESULTS AND DISCUSSION

### Construction of a VLP-based cGAMP delivery platform using bacteriophage MS2

Effective cellular delivery cGAMP requires addressing its susceptibility to enzymatic degradation as well as its membrane impermeability. We envisioned cGAMP delivery using a viral capsid delivery system based on the bacteriophage MS2 virus-like particle (MS2 VLP). The MS2 capsid consists of 180 identical coat proteins that self-assemble to form a hollow protein sphere 27 nm in diameter. Due to the sequence-defined nature of the MS2 coat protein, the internal and external surfaces can be separately engineered^41^. This has been leveraged to design a variant of MS2 with improved uptake in mammalian cells by adding two cationic mutations, T71K and G73R, on the exterior surface^39^. These residues are strategically positioned around 32 pores on MS2 where either 5 or 6 monomers converge; as a result, they create localized patches of positive supercharge that are understood to bind strongly to negatively charged heparan sulfate proteoglycans (HSPGs) on cell surfaces (**Figure 1a**, residues in blue). Importantly, this can be done without causing MS2 to become globally supercharged, as is the case for other polycationic carriers. As a result of this property, MS2 has been observed to avoid potential toxic effects, such as hemolysis and innate toxicity^39^. We observed this T71K/G73R MS2 mutant, referred to as KR MS2, to have vastly enhanced rates of cell uptake as well as delivery of otherwise membrane-impermeable small-molecule drug cargo^39,40^. In subsequent studies, we also observed that an S37P mutation showed modest improvements in delivery efficacy in some cell types, such as dendritic cells^42,43^. We proceeded with this mutant S37P/T71K/G73R mutant of MS2, which we will refer to as PKR MS2 herein.

**Figure 1.**
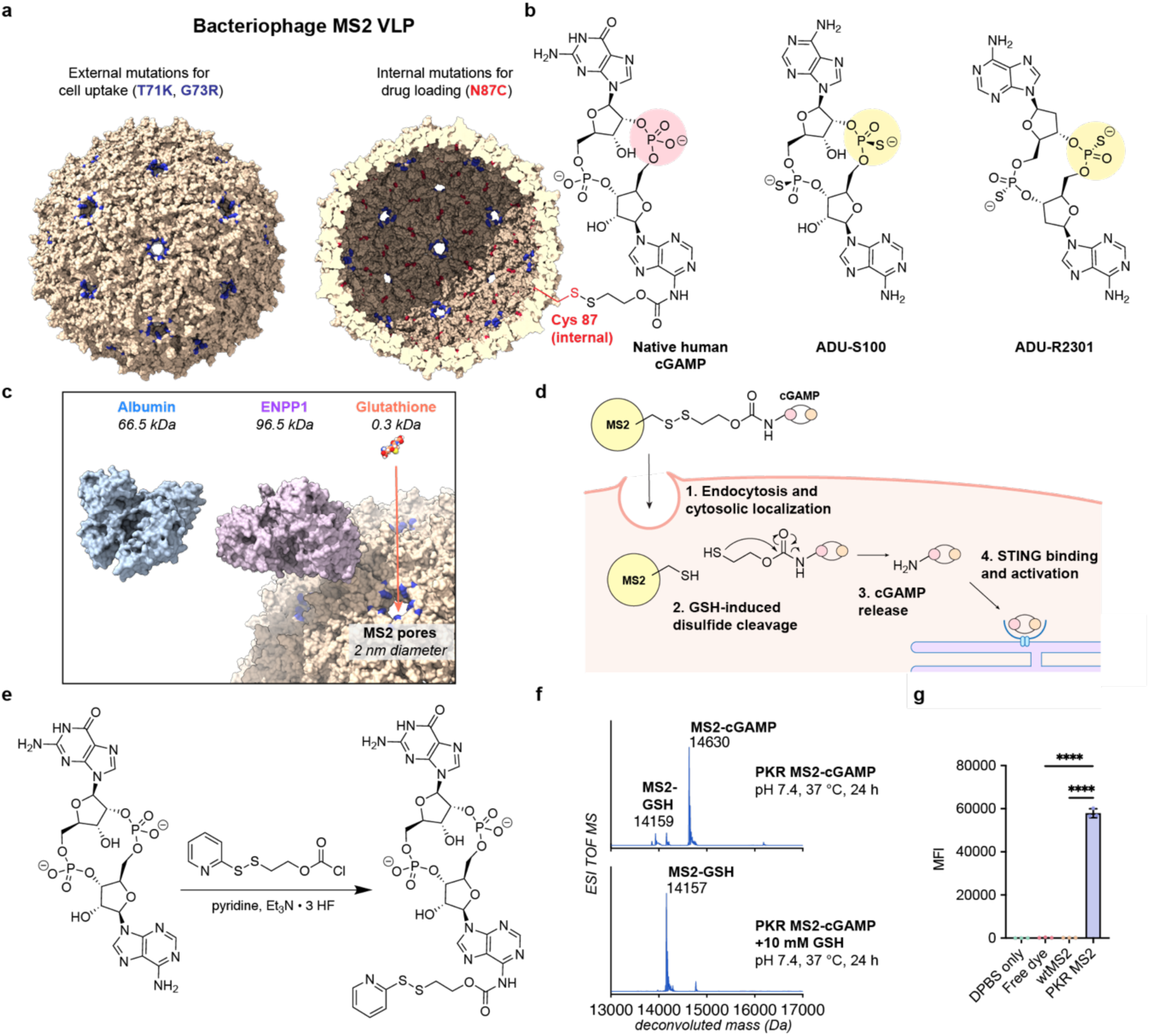
Design and construction of the MS2-cGAMP conjugate. **(a)** Structure of PKR MS2, showing external T71K/G73R mutations in blue and internal N87C mutations in red. cGAMP can be attached to the internal cysteine by a disulfide linker. **(b)** Comparison of native human cGAMP, bearing two phosphate groups, and two synthetic phosphorothioate CDN STING agonists. **(c)** The 2 nm pores of MS2 are too small to allow for the diffusion of proteins such as albumins or nucleases, but small molecules such as glutathione can pass through. **(d)** Diagram of disulfide cleavage and drug release in the cell. The disulfide attachment of cGAMP to the MS2 internal surface is stable in the extracellular environment; once internalized and localized to the reducing cytosolic environment, millimolar concentrations of glutathione diffuse into MS2 through its 2 nm pores and cleave the disulfide linker, which undergoes a self-immolative mechanism to release cGAMP in its native form. **(e)** Synthesis of disulfide linker-appended cGAMP. **(f)** ESI-QTOF mass spectra of PKR MS2 after cGAMP attachment, as well as treatment of the conjugate with 10 mM glutathione. **(g)** THP-1 monocyte cells were treated with fluorescently labeled PKR MS2 and controls, and cells were analyzed by flow cytometry to determine cell uptake.

As MS2 is not an enveloped virus, it cannot effectively retain small anionic cargo such as cGAMP simply through electrostatic interactions, as is the case with some other viruses^44,45^. However, by introducing a reactive cysteine residue on the capsid interior surface via a N87C mutation (**Figure 1a**), drugs and other cargo can be loaded inside the capsid using well-established thiol bioconjugation chemistries. The sequence-defined nature of the 180-mer MS2 capsid confers an advantage over enveloped and other native virus-based drug delivery carriers by ensuring a consistent and easily assayable drug concentration is loaded into each capsid. In the past, diverse cargoes such as paclitaxel^46^, doxorubicin^47^, monomethyl auristatins^39,43^, photodynamic therapy agents^48^, and peptide-based vaccines^42^ have been loaded into the capsid and delivered to cells with this platform.

Covalent loading techniques have been used by us and others in previous studies for synthetic CDN STING agonists, such as ADU-S100 and ADU-R2301, which generally bear the more nuclease-resistant phosphorothioate linkage (**Figure 1b**)^15,40,49^. As the phosphate group of native cGAMP is more prone to degradation, the delivery vehicle must not only provide the means to enter the cell and release cGAMP but also protect it from nucleases during transport.

To attach cGAMP to the interior of MS2, we envisioned using a reductively cleavable disulfide linker (**Figure 1a-b**), which we have previously found to be uniquely suited for drug loading in MS2^40^. The disulfide linker can be incorporated directly by modifying the interior Cys 87 residues, achieving up to 180 copies of drug per capsid. The interior of MS2 can be accessed by the 32 pores, each 2 nm in diameter, on its surface. This size allows small molecules such as cGAMP to diffuse in and out, while restricting proteins and larger molecules, such as soluble phosphatases that may degrade cGAMP^15,16^ or redox-sensitive serum albumins that may prematurely cleave the disulfide linker^50^ (**Figure 1c**). However, once inside the cell, a millimolar concentration of glutathione (GSH) and other small-molecule thiols^50^ can access the disulfide and cleave the drug from the capsid. A subsequent self-immolative mechanism ensures the traceless release of the drug (**Figure 1d**).

While most other studies have focused on synthetic phosphorothioate CDNs due to their higher serum stability, we can use native cGAMP (**Figure 1b**) due to the protective nature of MS2. This confers multiple potential advantages; as the evolved native agonist for STING, cGAMP would be expected to encounter fewer issues with toxicity, off-target effects, or unexpected metabolic behaviors in the body^51^. As a result of its rapid degradation, the transient nature of cGAMP as a signaling molecule could produce a more highly localized STING response that prevents problematically broad inflammation. Practically, cGAMP can also be accessed at scale more cheaply and conveniently than phosphorothioate CDNs through enzymatic synthesis from ATP and GTP using a recombinant version of the enzyme cGAS (**Figure S1a-b**)^52–54^.

Covalent attachment methods for synthetic CDNs generally use non-native functional groups as attachment sites; few reported methods exist for equivalent covalent modification of native cGAMP. Therefore, we sought to develop conditions to append a linker to cGAMP via the exocyclic amine of the CDN adenine base. This (**Figure 1e, Figure S1b**) was achieved via a variation of the protocol we used previously^40^, in which a pyridyl disulfide linker was attached to the adenosine base through a carbamate linkage. We found that for cGAMP, this reaction required the addition of triethylamine trihydrofluoride (Et_3_N • 3 HF) in pyridine for success. The purified product was then coupled to the internal cysteine on PKR MS2 through disulfide exchange, achieving high loading yields of 80-90%, corresponding to about 140-160 drug molecules per capsid (**Figure S1c**). Dynamic light scattering was used to ensure that cGAMP-loaded MS2 remained assembled in its ∼27 nm capsid form (**Figure S1d**). In addition, this disulfide linkage was confirmed to cleave and release cGAMP from MS2 in the presence of 10 mM glutathione over 24 h at 37 °C, while the conjugate remained intact in the absence of glutathione (**Figure 1f**).

### cGAMP-loaded PKR MS2 achieves more potent STING pathway activation compared to free cGAMP

We next evaluated the performance of cGAMP-loaded PKR MS2 capsids (PKR MS2-cGAMP) for the activation of STING pathway signaling in an *in vitro* cell model. This was done in a THP-1 monocyte cell model containing a CDN-inducible tdTomato reporter system that we previously established and demonstrated as an accurate proxy for STING activation and subsequent type I IFN response^19,40^. These cells can readily take up PKR MS2, as demonstrated by fluorescent dye-loaded capsids (**Figure 1g**). Upon 24 h treatment of the cells with STING-activating agents, PKR MS2 delivery of cGAMP showed a dramatic increase in potency compared to free cGAMP as defined by EC_50_ values. While free cGAMP exhibited an EC_50_ above 50 µM, the EC_50_ of PKR MS2-cGAMP dropped to 22 nM, representing a roughly 2000-fold improvement (**Figure 2a**). Even when compared to the drug ADU-S100, a synthetic CDN STING agonist bearing phosphatase-resistant phosphorothioate groups, PKR MS2-cGAMP still exhibited a greater than 400-fold increase in potency (**Figure 2a**), suggesting that PKR MS2 enhances cGAMP activity not only through protection of the cargo but also an enhanced delivery of the drug to the cell cytosol. STING activation efficacy, through monitoring of IFN-beta mRNA, was also confirmed by RT-qPCR measurements of the reporter cells (**Figure 2b**).

**Figure 2.**
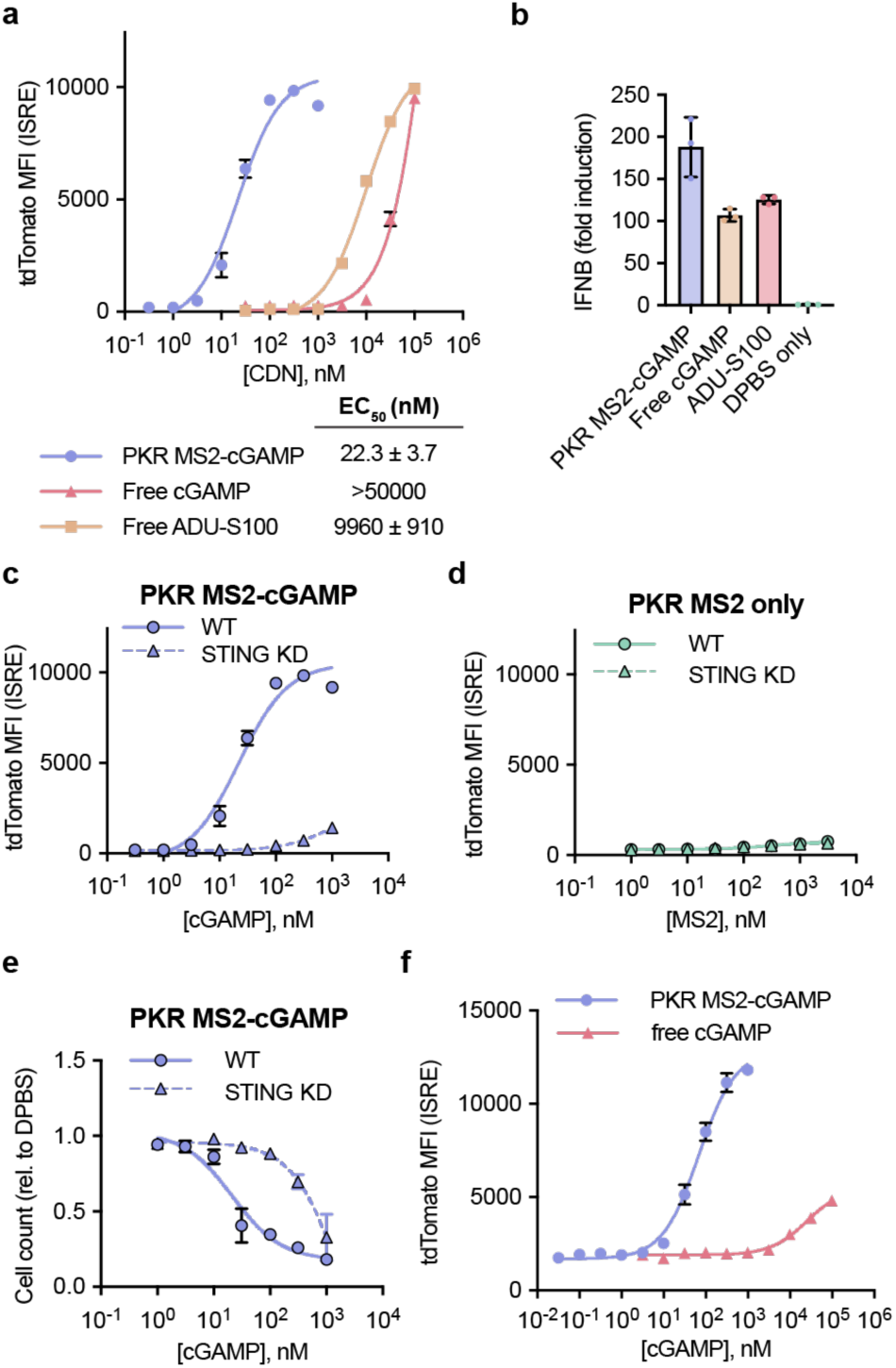
STING activation by PKR MS2-cGAMP. **(a)** THP-1 monocyte cells containing a tdTomato STING reporter system were treated with PKR MS2-cGAMP, free cGAMP, and free ADU-S100 for 24 h and analyzed by flow cytometry. Median fluorescence intensity values were plotted to calculate EC_50_ values. **(b)** THP-1 reporter cells were treated with 100 nM PKR MS2-cGAMP, 100 µM free cGAMP, or 10 µM free ADU-S100 for 24 h, and the IFN-beta mRNA (IFNB1) was isolated and analyzed by RT-qPCR. **(c, d)** Type I IFN response in THP-1 reporter cells with wild-type STING (WT) was compared to that in reporter cells in which STING expression was knocked down by transduction with an shRNA construct (STING KD). **(e)** STING wild-type and STING knockdown THP-1 reporter cells were treated with PKR MS2-cGAMP for 24 h and compared for cell viability by MTS assay. **(f)** THP-1 reporter cells differentiated into macrophages using PMA were treated with PKR MS2-cGAMP or free cGAMP for 24 h and analyzed by flow cytometry.

To ensure that the principal cause of the type I IFN response was from STING activation rather than any potential activity of the MS2 itself, the interferon reporter activity in wild-type THP-1 reporter cells was compared to that in a STING knockdown (STING KD) version of the same cell line^55^. For all treatments, STING KD reporter cells elicited little-to-no interferon response at high treatment concentrations (**Figure 2c-d, Figure S2a-b**), indicating that the interferon responses from PKR MS2-cGAMP, free cGAMP, and ADU-S100 cells were all from STING activation. Additionally, PKR MS2 alone elicited no activity in STING WT or STING KD reporter cells (**Figure 2d**). At the highest concentrations of PKR MS2-delivered cGAMP, a minor reporter signal was observed with STING KD cells, which was likely due to a combination of incomplete ablation of STING expression in the knockdown cell line and potential activation of toll-like receptors from trace impurities of lipopolysaccharide or RNA left over from the production of PKR MS2 in *E. coli*.

To ensure that PKR MS2 does not cause any unwanted toxic effects as a delivery agent, the cell viability of THP-1 reporter cells was observed after 24 h treatment (**Figure 2e**). The viability of PKR MS2-cGAMP treated cells closely followed the STING activation curve, which is to be expected since STING activation can lead to various cell death mechanisms in monocytes and other cell types^56,57^. In contrast, PKR MS2-cGAMP in STING KD cells did not exhibit decreased cell viability except at the highest tested concentrations, confirming that cell death was principally from STING-dependent processes rather than any systemic toxicity from the drug formulation.

STING activity was also observed in the same reporter cells differentiated into macrophages using phorbol 12-myristate 13-acetate (PMA). Again, in those cells, STING activity was greatly enhanced through PKR MS2 delivery of cGAMP (**Figure 2f**). This was consistent with previous reports finding effective delivery to macrophages^40^, as well as data that support the ability of MS2 capsids bearing the T71K/G73R mutations to be taken up by a broad range of cell types^39,42^.

### PKR MS2 protects cGAMP from degradation outside the cell

The use of PKR MS2 to deliver cGAMP to the cell cytosol highlights its advantages over using both free cGAMP as well as other drug design or delivery approaches. One of the key limitations of native cGAMP as a therapeutic agent is its susceptibility to endogenous nucleases. The most well-studied of these, ectonucleotide pyrophosphatase/ phosphodiesterase 1 (ENPP1), is a negative regulator of STING signaling^15^. ENPP1 is expressed ubiquitously and either anchored to the cell exterior surface or secreted into the extracellular space, where it can degrade extracellular cGAMP as it is released from cells^16^. It is known that ENPP1 is upregulated in some cancers, which can suppress STING signaling and lead to poorer prognoses^58,59^. Conversely, small-molecule inhibitors of ENPP1 have indirectly led to increased STING signaling and immune response through decreased cGAMP degradation^59,60^. That being said, extracellular degradation by ENPP1 and other less well-characterized native enzymes is a major impediment in administrating cGAMP therapeutically.

Protected inside the MS2 capsid, cGAMP is inaccessible to these soluble or membrane-bound nuclease enzymes that are too large to diffuse through the 2 nm MS2 pores (**Figure 1c**). We demonstrated this by comparing STING activation by PKR MS2-cGAMP, free cGAMP, or ADU-S100 in the presence and absence of non-heat inactivated fetal bovine serum (FBS), which is known to contain soluble cGAMP-degrading phosphatases^15^. STING activity was also analyzed in the presence of serum as well as STF-1084, an inhibitor of ENPP1 that is known to improve STING activation elicited by cGAMP^60^. The nanomolar STING activity of PKR MS2-cGAMP was largely maintained with or without serum, as well as in the presence of STF-1084 (**Figure 3a**). Conversely, for free cGAMP, addition of serum caused a roughly 3- to 10-fold decrease in STING activation (**Figure 3b**). This loss of activity was partially but not completely recovered by STF-1084, which could hint that other serum enzymes may be involved in cGAMP clearance. Furthermore, an equivalently large decrease in activity was observed for the phosphorothioate drug ADU-S100, despite its known resistance to traditional phosphatases, and unlike native cGAMP, ADU-S100 activity was not recovered by STF-1084 (**Figure 3c**). These results highlight the challenges of identifying and inhibiting all possible extracellular degradation pathways of free CDN drugs. Even the synthetic CDNs that bear phosphorothioates, hailed as nuclease-resistant versions of traditional phosphates in nucleotide drugs, only exhibit reported serum half-lives on the order of a few hours^61^, and the method by which they are cleared is not well understood. Designed to protect against all enzymatic degradation mechanisms, PKR MS2 provided far superior protection to cGAMP in these experiments.

**Figure 3.**
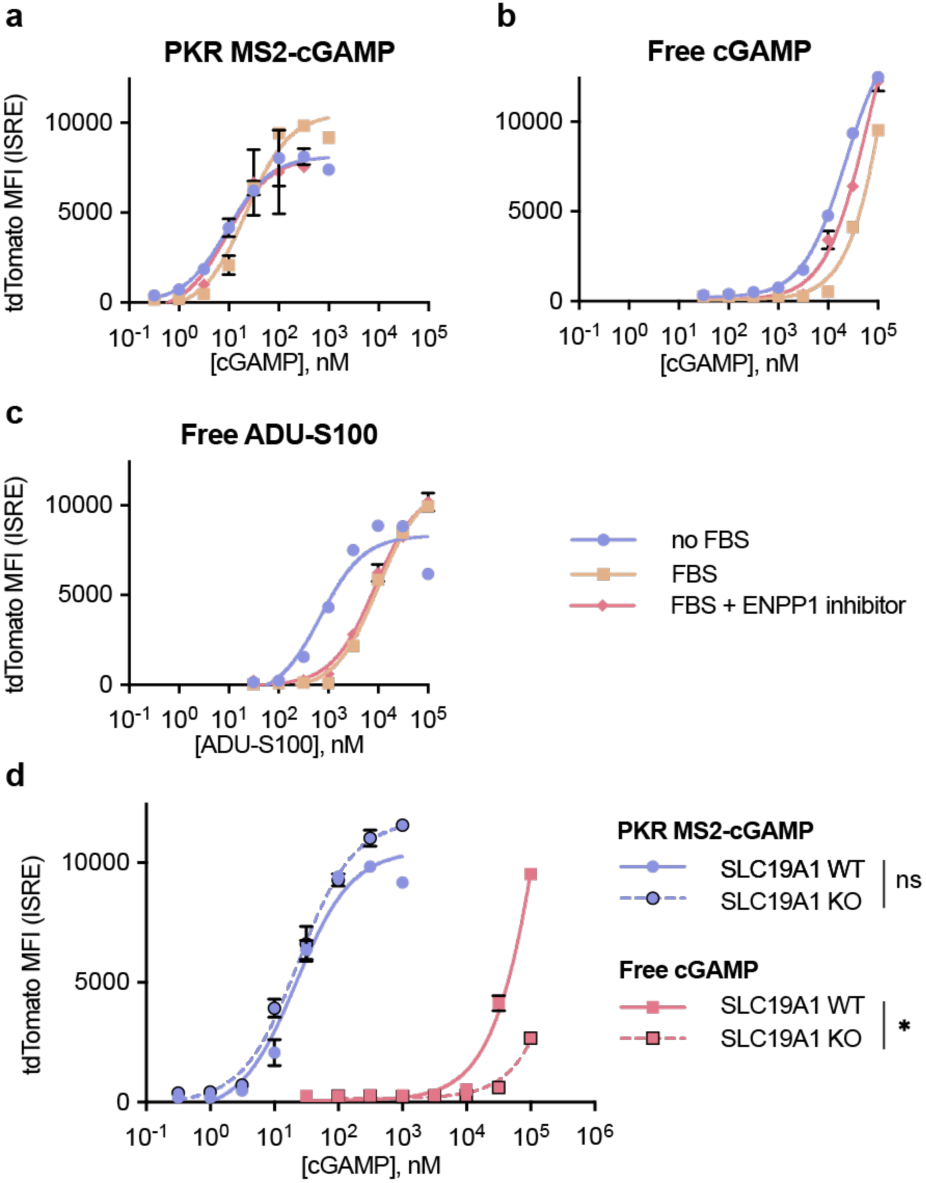
Mechanism of cGAMP protection and uptake by MS2. **(a-c)** THP-1 reporter cells were treated with PKR MS2-cGAMP, free cGAMP, or free ADU-S100 in the absence of nuclease-containing fetal bovine serum (FBS) or the presence of ENPP1 inhibitor STF-1084 and compared to the data from Figure 2a. **(d)** THP-1 reporter cells with and without a knockout of the cell-surface transporter SLC19A1 were treated with PKR MS2-cGAMP and free cGAMP for 24 h and compared.

### PKR MS2 delivers cGAMP across the cell membrane independent of cell-surface transporters

As native cGAMP and synthetic small-molecule CDNs are impermeable to the cell membrane due to their anionic charge, they must rely on small-molecule anion transporter proteins on cell surfaces for cytosolic entry. These transporters, which include SLC19A1^19,52^, SLC46A2^62^, and others^20,63^, have relatively low affinity for CDNs, can be inconsistently expressed across cell types, and have the potential to be downregulated in certain cancers^59^. PKR MS2, which enters the cell through endocytic means^40^, is not expected to depend on these transporters. To demonstrate this, STING activity in THP-1 reporter cells was compared to that in an analogous THP-1 reporter cell line containing a knockout for SLC19A1, an anion transporter that serves as the major cGAMP transporter in THP-1 cells (**Figure 3d**)^19,52^. While free cGAMP lost much of its already-weak activity in the absence of the SLC19A1 transporter, the activity of PKR MS2-cGAMP was not affected at all, confirming that it operates independently of SLC19A1 and related cell-surface transporters.

Previous data exploring the uptake of MS2 capsids bearing the T71K/G73R mutations revealed that MS2 capsids have high affinity to heparan sulfate groups on cell surfaces, which cause binding to cells and subsequent endocytosis^39^. This uptake mechanism, as well as that of the subsequent delivery of cGAMP, was investigated in this study in THP-1 monocytes. PKR MS2 attached to a fluorescent dye was delivered to cells in the presence of a number of potential inhibitors to reveal the mechanism of uptake and drug delivery, respectively. The inhibitors used were soluble heparin to reduce MS2 binding to cells, cytochalasin D to inhibit phagocytosis, dynasore to inhibit clathrin-mediated endocytosis, and chloroquine to inhibit lysosomal acidification^39,64^ (**Figure S2b**). While heparin eliminated uptake into cells, all other endocytosis inhibitors showed insignificant changes. These results support previous findings that the endocytic pathway of MS2 uptake may occur through multiple redundant mechanisms and that heparan sulfate binding is a key part of the process.

To probe the mechanism of cGAMP delivery to the cytosol post-endocytosis, PKR MS2-cGAMP and free cGAMP activity was compared in the absence or presence of chloroquine. After endocytosis, endosomes undergo acidification and subsequent fusion to lysosomes, a process inhibited by chloroquine, which accumulates in endosomes and acts as a neutral pH buffer^65^. Chloroquine was observed to cause an enhancement of STING signaling by free cGAMP, a known effect due to reduced autophagy of the STING receptor and other signaling proteins (**Figure S2c**)^66^. However, addition of chloroquine to PKR MS2-cGAMP elicited a modest *reduction* in STING signaling of roughly 4-fold (**Figure S2c**). This suggests that endosomal acidification and lysosomal processing facilitate the release of cGAMP from PKR MS2 at least to some degree. While work is ongoing to characterize the behavior of MS2 inside the endosome in more detail with the goal of maximizing cytosolic drug release, it remains the case that MS2 is able to release enough of its cGAMP cargo within cells to access the cytosolic facing STING receptor and achieve large improvements in STING activation compared to free cGAMP.

### PKR MS2-cGAMP shows enhanced STING-dependent *in vivo* antitumor effects in a murine colon carcinoma model

Having established the enhanced efficacy of PKR MS2-cGAMP at STING signaling activation in cell culture, we next examined its potential for antitumor responses *in vivo*. As discussed previously, STING signaling in a tumor microenvironment can activate lymphocytes and antigen-presenting cells, such as macrophages and dendritic cells, leading to potent and long-lasting antitumor immune responses^4–8^. We aimed to compare the activity of PKR MS2-cGAMP to both free cGAMP and ADU-S100 in a tumor model. The MC38 colon carcinoma model, compatible in syngeneic immunocompetent C57BL/6J mice, was chosen due to its known response to STING agonists in previous studies^5,36,38^. PKR MS2 was observed to enter MC38 cells (**Figure S2d**), but the same cells treated with STING agonists did not exhibit any upregulation in STING-dependent type I IFN signaling (**Figure 4a**), suggesting a deficiency in STING signaling by the tumor cells themselves and signifying that any STING-dependent antitumor immune response would require activation of STING signaling from host immune cells in an *in vivo* system.

**Figure 4.**
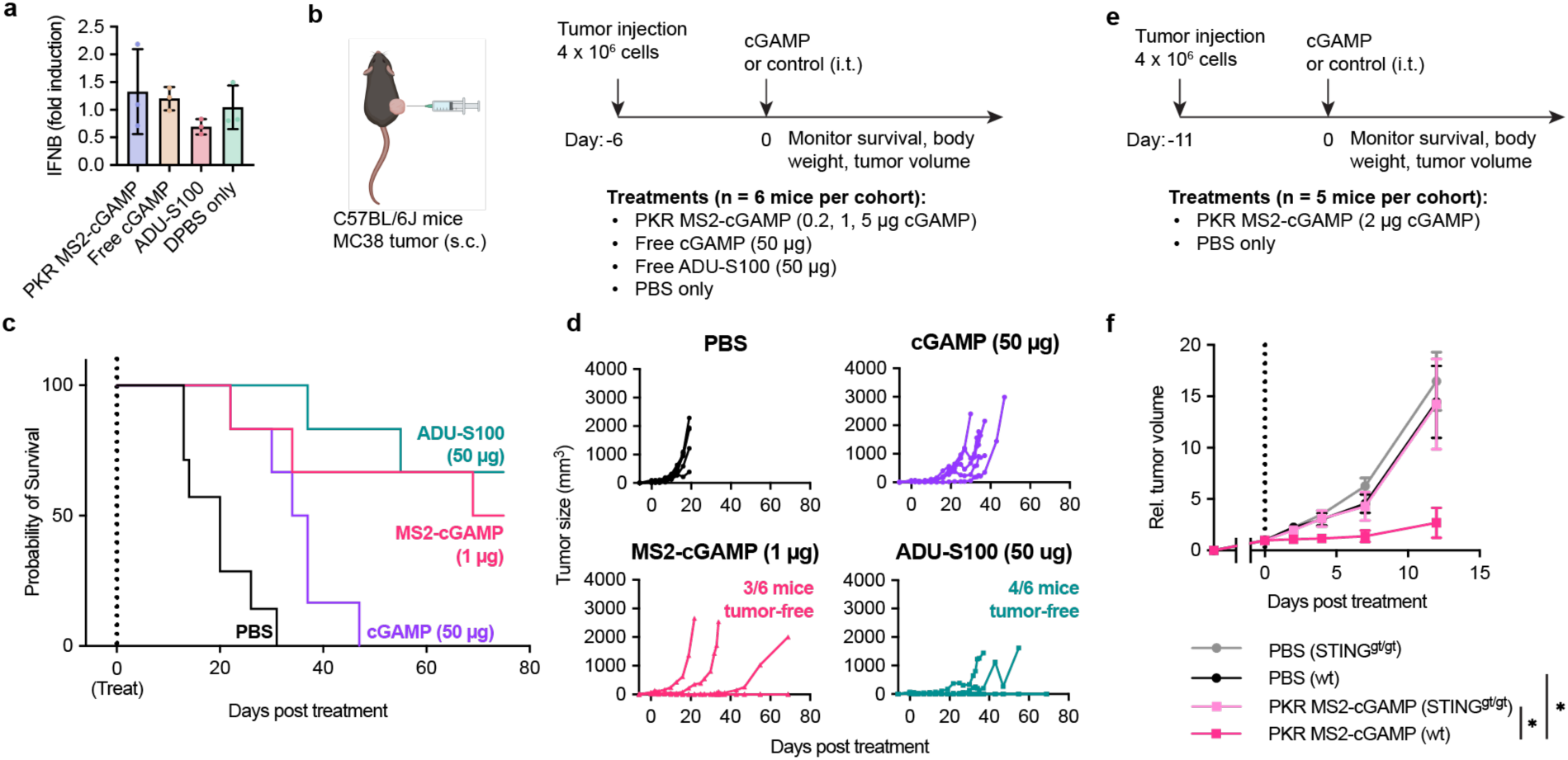
*In vivo* antitumor efficacy of PKR MS2-cGAMP in a murine colon carcinoma model. **(a)** MC38 colon carcinoma cells were treated with PKR MS2-cGAMP, free cGAMP, or ADU-S100 for 24 h and analyzed for IFN-beta mRNA levels by RT-qPCR. **(b)** Experimental design for subcutaneous tumor injection and treatment in mice. **(c)** Kaplan-Meier survival curves of the lowest effective dosage of PKR MS2-cGAMP and free drug controls. **(d)** Individual tumor volumes in each treatment arm over time. **(e)** Experimental design for antitumor comparison in STING^gt/gt^ versus wild-type (wt) mice. Mice were injected with MC38 tumor cells and subsequently treated with PKR MS2-cGAMP or a PBS control. **(f)** Relative tumor volumes of wt and STING^gt/gt^ mice after PKR MS2-cGAMP or PBS treatment. Tumor volumes were normalized to the average tumor volume of each cohort on treatment day (Day 0), and the resulting curves were compared through day 12 (prior to any animal deaths in control arms) by ANOVA.

It has been established that a single 50 µg dose of ADU-S100 injected intratumorally can generate an effective antitumor response in the *in vivo* MC38 model^22^; an equivalent mass of native cGAMP was used as a control as well. The *in vivo* activity of PKR MS2-cGAMP was not known; given its *in vitro* potency, three dosage levels were tested, representing 5 µg, 1 µg, and 0.2 µg cGAMP, as defined by the mass of the cGAMP component of the drug. MC38 tumors were first established subcutaneously into C57BL/6J mice, followed by administration of each STING agonist treatment or DPBS control via intratumoral injection. Groups of n = 6 mice were included in each cohort. Mice were then monitored for survival and tumor volumes (**Figure 4b**).

Mice treated with a single dose of PKR MS2-cGAMP down to 1 µg drug were observed to have comparable survival to those receiving the standard 50 µg dosage of free ADU-S100 (**Figure 4c**). In contrast, mice receiving a 50 µg dose of free cGAMP showed only modest long-term survival increases. In observing tumor volumes, 1 µg of PKR MS2-cGAMP was able to reduce and completely eliminate tumors in half the mice in the treatment cohort, comparable to the efficacy of 50 times that amount of ADU-S100. In contrast, free cGAMP was able to slow tumor growth but not reduce tumor size, even at the 50-fold higher dosage (**Figure 4d**). Additional PKR MS2-cGAMP at a 5 µg dosage did not have any added antitumor effect, and the lower 0.2 µg dosage failed to induce a significant antitumor response (**Figure S3a-b**). Furthermore, monitoring of initial weight loss in mice after drug treatment showed that 1 µg PKR MS2-cGAMP caused a comparable weight loss as 50 µg ADU-S100, while 5 µg PKR MS2-cGAMP caused increased weight loss (**Figure S3c**). Using weight loss as a proxy for systemic toxicity effects, these results established an optimal dosage window for PKR MS2-cGAMP in mice to be around 1 µg, balancing maximal antitumor efficacy with minimal off-target toxicity.

While the most likely cause of the antitumor effect was a STING-based immune system response, we wanted to rule out a localized cytotoxic response in the tumor cells alone or a different immunogenic response to the MS2 capsids. To do this, a similar study was conducted comparing the responses of the same MC38 tumor cells in STING^gt/gt^ (STING-deficient) mice, which harbor a single nucleotide point mutation in the *Sting* gene that causes it to act as a null allele that fails to produce STING protein^55^, versus wild-type (WT) C57BL/6J mice with fully functional STING (**Figure 4e**). Tumor volumes were compared among the cohorts during the initial two weeks of the experiment prior to animal morbidity requiring euthanasia in control groups. While PKR MS2-cGAMP treatment again elicited tumor reduction in WT mice, this antitumor effect was not observed in tumor-bearing STING-deficient mice (**Figure 4f**). A more prolonged survival was also observed with PKR MS2-cGAMP in WT mice, albeit to a less significant extent (**Figure S3d-e**); this was likely because the tumors in STING^gt/gt^ mice happened to be modestly smaller on the day of treatment than those in WT mice (**Table S2**). A slight survival increase for STING^gt/gt^ mice treated with PKR MS2-cGAMP could be attributed to weaker toll-like receptor or other STING-independent immune responses to the capsid formulation, such as those that were observed in the *in vitro* studies in STING KD cells (**Figure 2c**). Nonetheless, these data demonstrate that the antitumor effect of PKR MS2-cGAMP is STING dependent.

In summary, delivery of native cGAMP by PKR MS2 to an *in vivo* colon carcinoma model successfully achieved a STING-dependent antitumor response at a 50-fold lower dosage compared to the free synthetic CDN ADU-S100, demonstrating enhanced drug delivery to cells of interest. No such antitumor response was achievable with free cGAMP, highlighting the ability of PKR MS2 to protect its cGAMP cargo from degradation and clearance as well as facilitate its cell uptake and release to reach the cytosol. These results demonstrate the first instance of effective drug delivery using cationic KR mutation-bearing MS2 viral capsids to an *in vivo* tumor model, as well as the most efficient tumor treatment study using a single dose of native cGAMP reported to date.

## CONCLUSION

In this study, we demonstrated the use of a modified virus-like particle, bacteriophage MS2, as a delivery agent for the native STING agonist molecule cGAMP. By covalently attaching cGAMP to the occluded capsid interior to protect it from degradation and providing a method to facilitate its cell uptake and cytosolic release, PKR MS2 achieved a roughly 2000-fold enhanced potency for STING activation by cGAMP in an *in vitro* cell model. PKR MS2-cGAMP also demonstrated an effective STING-driven antitumor immune response *in vivo* at a 50-fold lower dosage than free cGAMP and ADU-S100, a leading synthetic cyclic dinucleotide STING agonist. This single low dose of PKR MS2-delivered cGAMP yielded curative antitumor effects in half of the treated mice, which could not be achieved even with a high dose of free cGAMP.

While cGAMP has been used in conjunction with other nanoscale drug delivery vehicles, these results uniquely demonstrate the use of a covalent linker to attach and release native cGAMP without any modification scars, ensuring that binding to the STING receptor is not compromised. The high cell uptake and internalization rate of PKR MS2 also resulted in a superior enhancement in STING activation compared to other delivery platforms. While it is not clear whether native cGAMP has any advantages over synthetic CDN or non-nucleotide STING agonists, the role of 2’,3’-cGAMP in cell signaling is not fully understood and remains an area of active research.

The therapeutic usage of CDNs for STING activation has shown repeated promise in preclinical murine models but has yielded consistently disappointing outcomes in clinical trials^9,12,23^. The inherent un-“drug-like” nature and resulting poor bioavailability of cGAMP complicates the rational design of STING agonists, and many lead compounds require impracticably high doses for therapeutic responses^12^. This poor uptake into relevant human cells is likely at least partially responsible for the limited clinical efficacy of these CDN drugs. Therefore, the ability to deliver cGAMP and achieve antitumor activity at much lower dosages and with only a single treatment could lead to greatly improved cellular uptake and stronger immunogenic responses in human subjects.

Beyond cGAMP, many native signaling molecules are known to bind and modulate disease-relevant receptors, enzymes, or other signaling proteins. While each of these molecules could themselves act as potential therapeutics, their limited cell uptake, unfavorable bioavailability, and serum instability often cause such bio-mimics to be discounted early in rational drug design in favor of more “drug-like” synthetic lead compounds that require extensive optimization campaigns and frequently have undesired off-target effects. The capsid-based delivery strategy, successfully demonstrated here with cGAMP, can serve as a blueprint for enabling the use of other such native signaling molecules with poor pharmacokinetics for therapeutic effect.

## AUTHOR INFORMATION

### Author Contributions

The manuscript was written through contributions of all authors. All authors have given approval to the final version of the manuscript.

### Notes

The authors declare no conflicts of interest.

## ACKNOWLEDGMENT

This work was supported by the Panattoni Family as well as the National Institutes of Health grants R01CA270790 and R01CA303968 (to D.H.R.). H.S.M. was supported by the National Science Foundation Graduate Research Fellowship Program (DGE 2146752). Any opinions, findings, conclusions, or recommendations expressed in this material are those of the authors and do not necessarily reflect the views of the National Science Foundation. Y.J. was supported by the Postdoctoral CIRM Training Program (EDUC4-12790) and the Cancer Research Institute / Amgen Irvington Postdoctoral Fellowship (CRI3984).

For Table of Contents only:

---

For Table of Contents only:

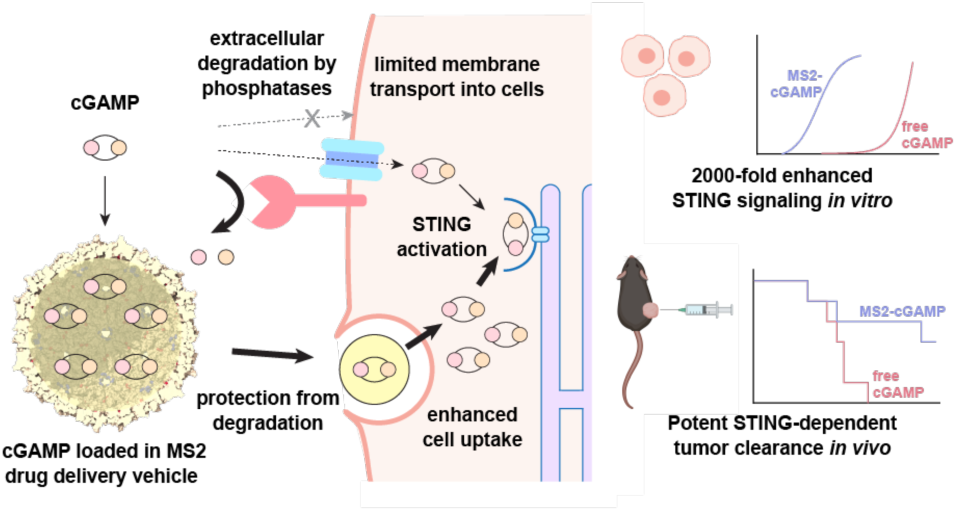

**Figure S1.**
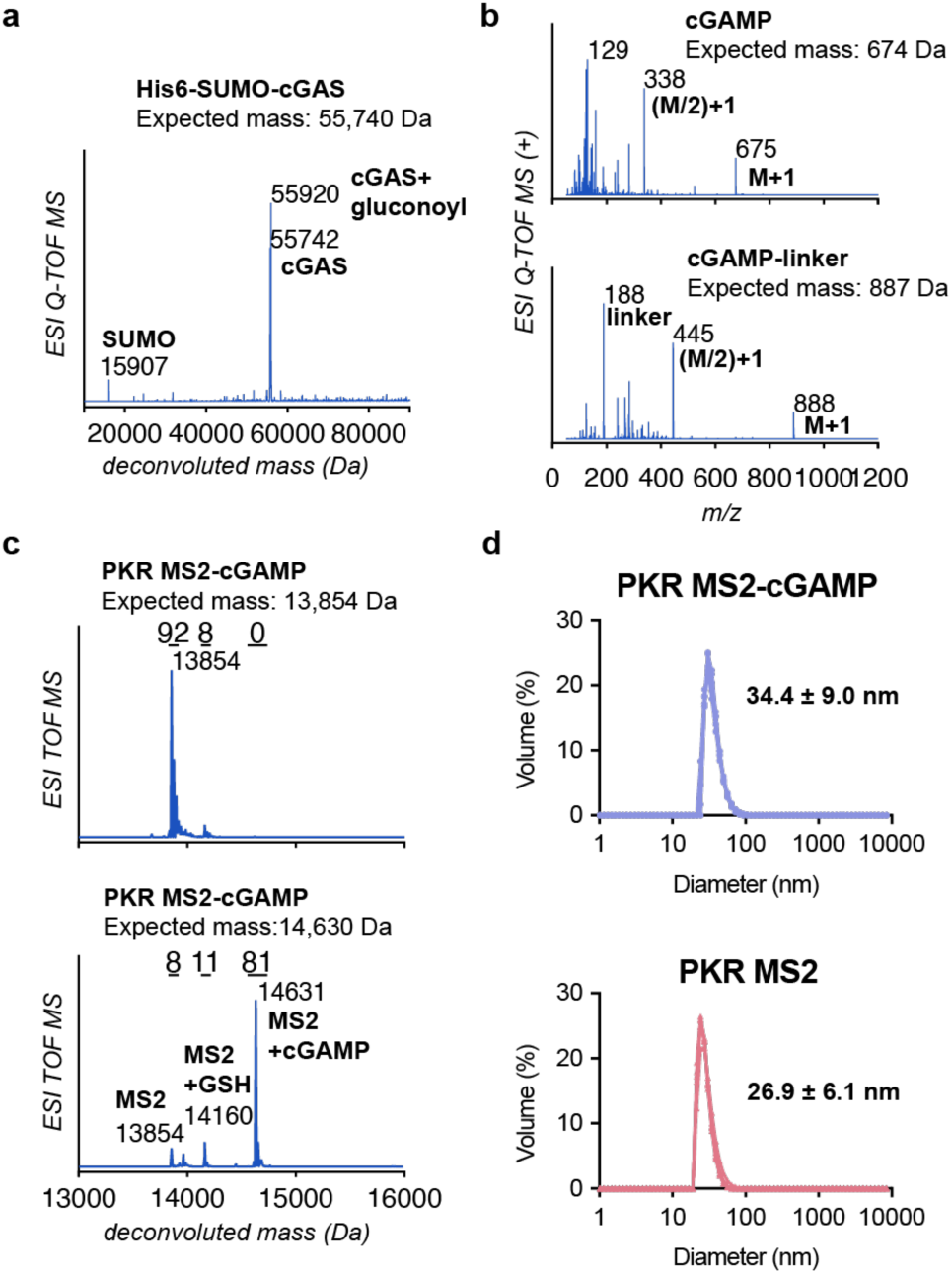
Construction and assembly of PKR MS2-cGAMP and components. **(a)** LC/MS spectrum of human truncated cGAS protein, expressed with a hexahistidine-SUMO tag for ease of purification. **(b)** LC/MS spectra of 2’,3’-cGAMP produced enzymatically as well as cGAMP-disulfide conjugate **2** (from **Scheme S1**) produced synthetically. **(c)** LC/MS spectra of PKR MS2 before and after conjugation to cGAMP. **(d)** Dynamic light scattering (DLS) spectra of PKR MS2 before and after attachment of cGAMP.

**Figure S2.**
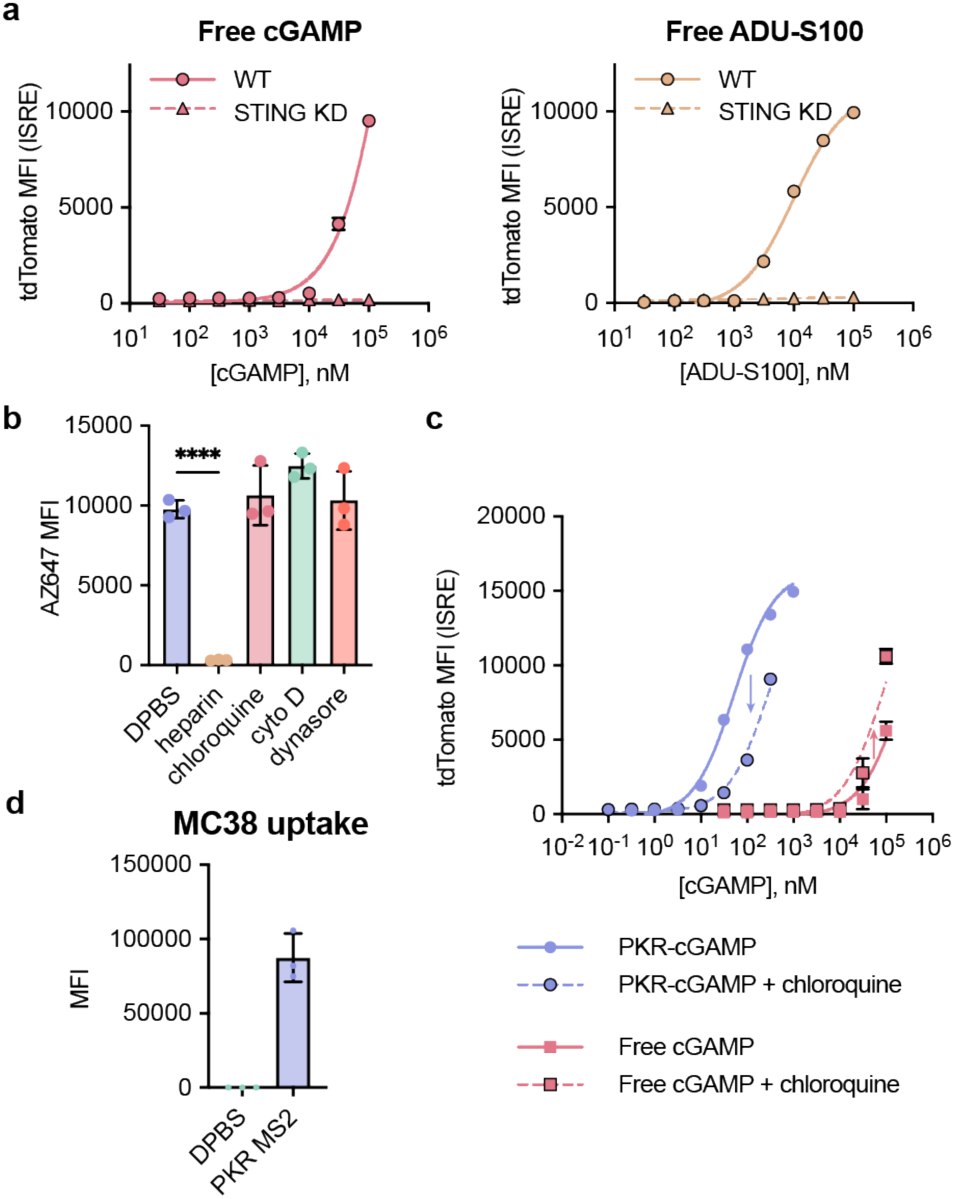
*In vitro* uptake and STING activation effects. **(a)** Type I IFN response in THP-1 reporter cells with wild-type STING (WT) was compared to that in STING knockdown (STING KD) reporter cells. **(b)** THP-1 STING reporter cells were treated with 200 nM PKR MS2 conjugated to a fluorescent AZFluor 647 maleimide dye in the presence or absence of various uptake and endocytosis inhibitors. Cells were analyzed for uptake by flow cytometry. **(c)** THP-1 STING reporter cells were treated with PKR MS2-cGAMP or free cGAMP in the presence or absence of 50 µM of the endosomal acidification agent chloroquine. Cells were analyzed for STING activation by flow cytometry. **(d)** MC38 cells were treated with 200 nM PKR MS2-dye conjugate or control and analyzed by flow cytometry for uptake.

**Figure S3.**
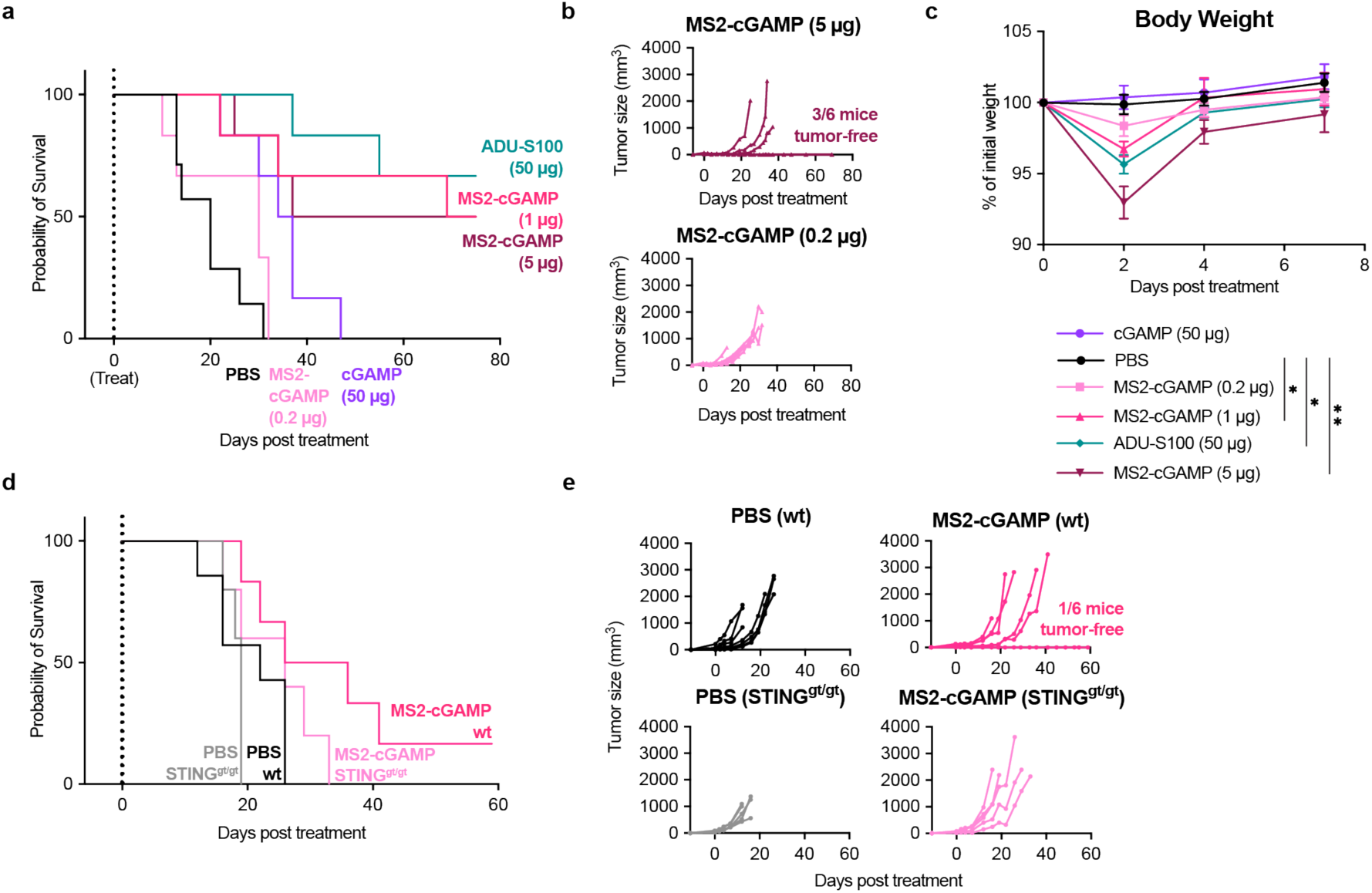
*In vivo* antitumor efficacy of PKR MS2-cGAMP. **(a)** Kaplan-Meier survival curves for mice treated with a single injection of 0.2 µg and 5 µg PKR MS2-cGAMP. Curves for mice cohorts shown in Figure 4c are also shown for comparison. **(b)** Individual tumor volumes over time for 0.2 µg and 5 µg PKR MS2-cGAMP dosage cohorts. **(c)** Relative body weight of all mice treated with PKR MS2-cGAMP, free cGAMP, free ADU-S100, or PBS control. Body weights were compared by ANOVA, with the statistical analysis shown representing significant body weight changes in treatment arms versus PBS control mice on day 2 after treatment. Mice in all treatment arms recovered their body weight within 7 days. **(d)** Kaplan-Meier survival curves for wt C57BL/6J and STING^gt/gt^ mice treated with PKR MS2-cGAMP or PBS control. **(e)** Individual tumor volumes over time for wt C57BL/6J and STING^gt/gt^ mice treated with PKR MS2-cGAMP or PBS control.

**Table S1.**
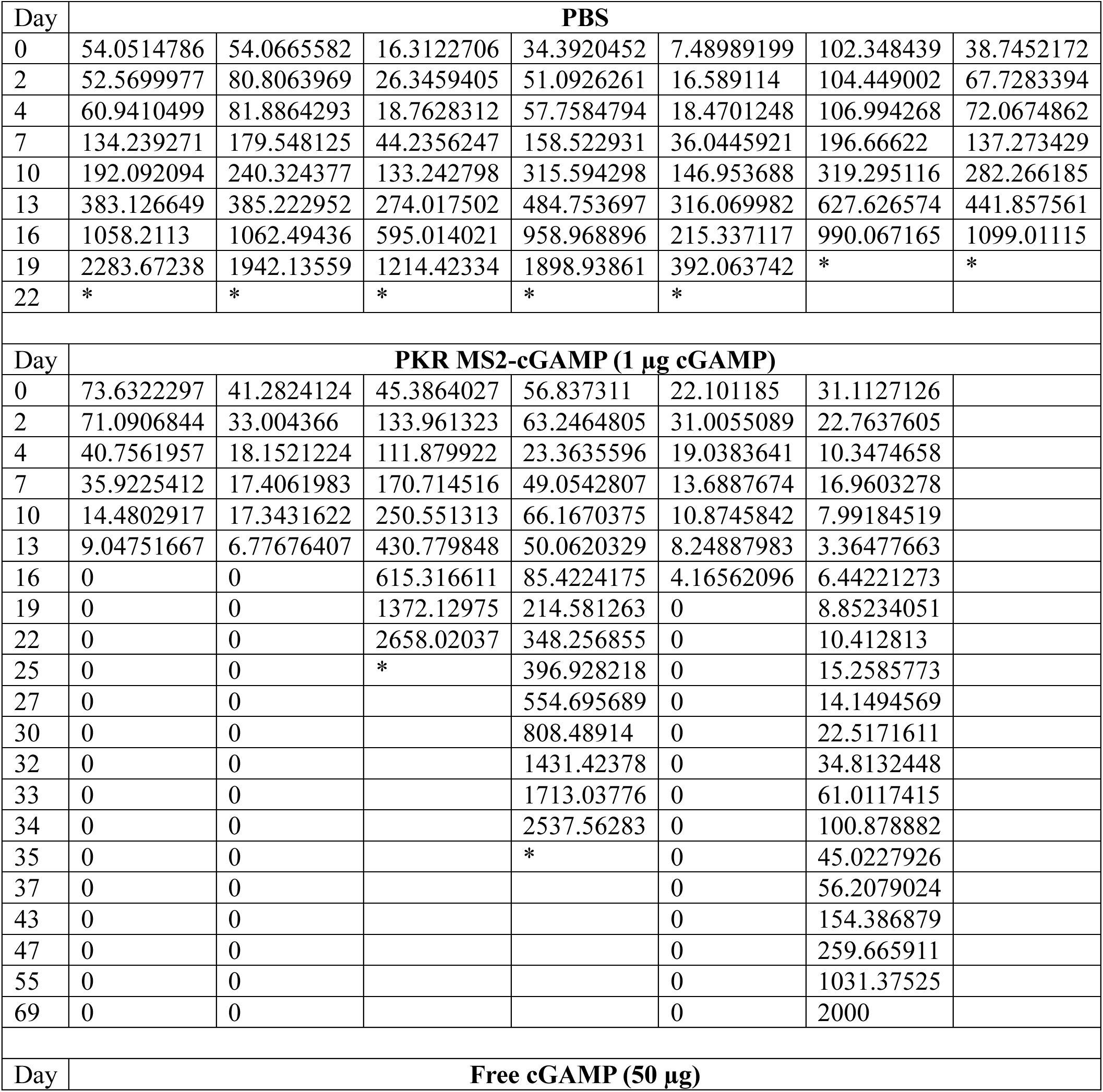
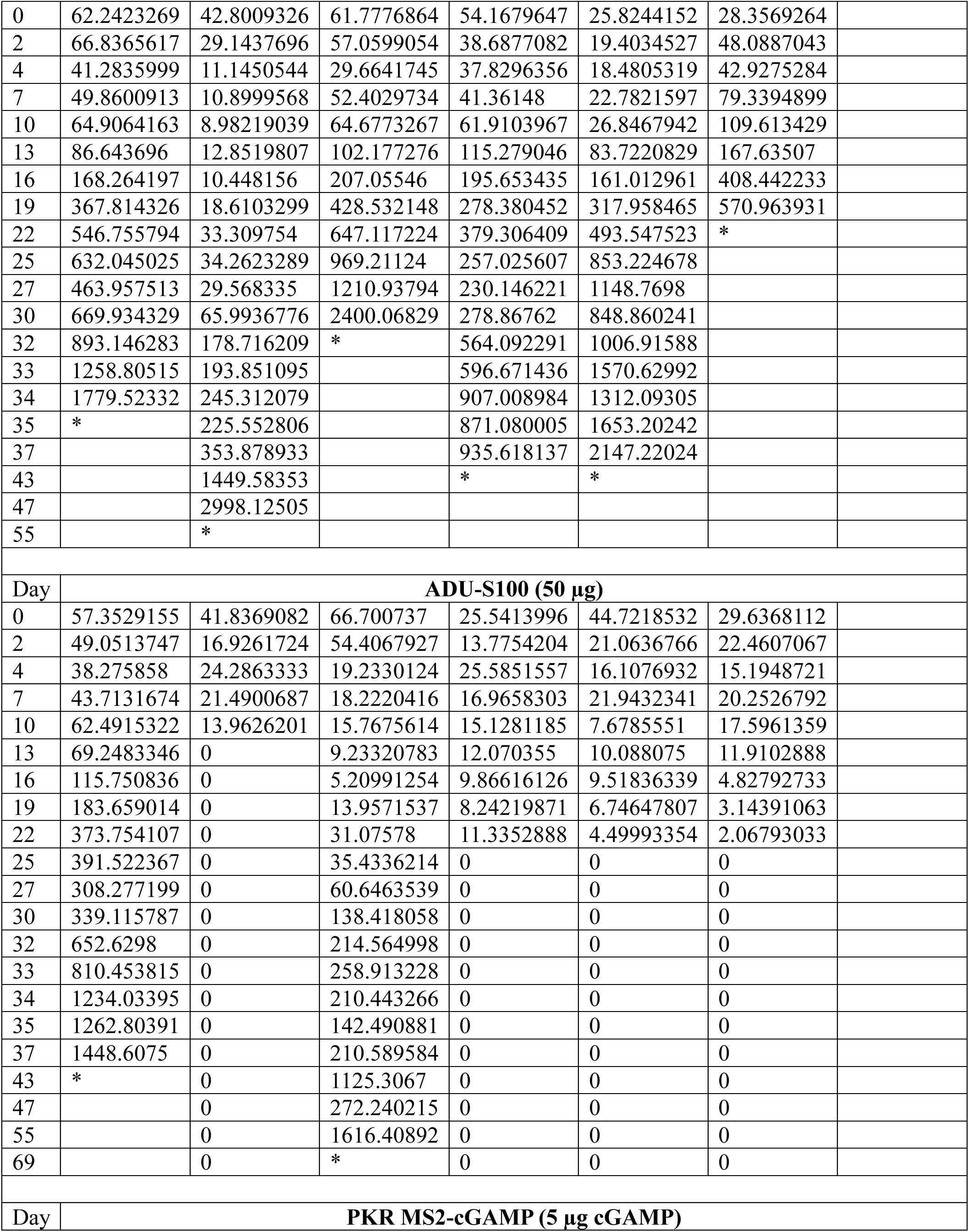
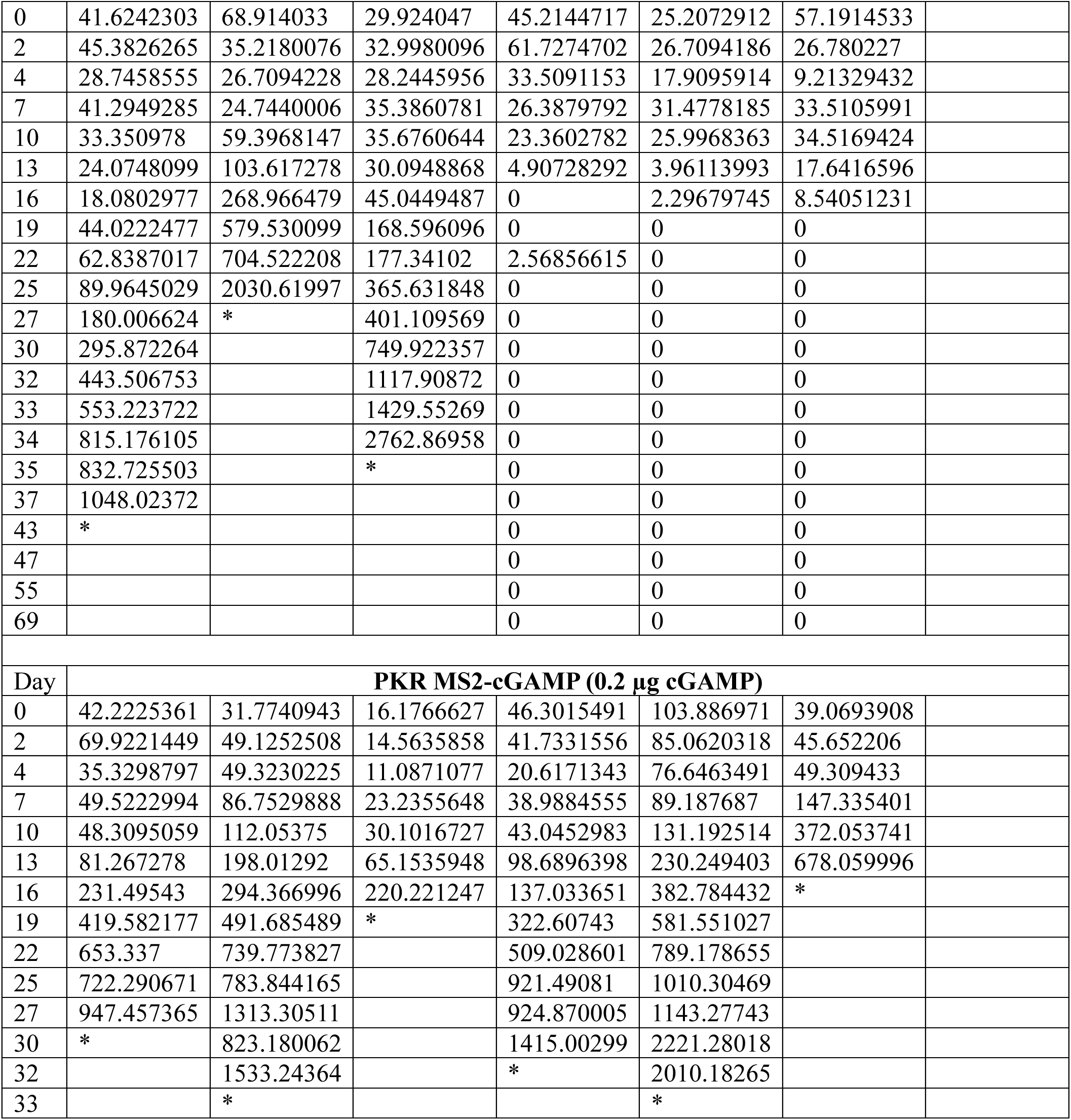
Tumor volumes in mice from the *in vivo* tumor response study represented in **Figure 4b**. All values are given in mm^3^. Each column represents the tumor volumes of one mouse in each sample. Mice that were sacrificed or died during the study are marked with an asterisk (*), while mice with no measurable tumor volume are indicated with a zero (0). The study was concluded 69 days after STING agonist or control treatment.

| Day | PBS |  |  |  |  |  |  |
| --- | --- | --- | --- | --- | --- | --- | --- |
| 0 | 54.0514786 | 54.0665582 | 16.3122706 | 34.3920452 | 7.48989199 | 102.348439 | 38.7452172 |
| 2 | 52.5699977 | 80.8063969 | 26.3459405 | 51.0926261 | 16.589114 | 104.449002 | 67.7283394 |
| 4 | 60.9410499 | 81.8864293 | 18.7628312 | 57.7584794 | 18.4701248 | 106.994268 | 72.0674862 |
| 7 | 134.239271 | 179.548125 | 44.2356247 | 158.522931 | 36.0445921 | 196.66622 | 137.273429 |
| 10 | 192.092094 | 240.324377 | 133.242798 | 315.594298 | 146.953688 | 319.295116 | 282.266185 |
| 13 | 383.126649 | 385.222952 | 274.017502 | 484.753697 | 316.069982 | 627.626574 | 441.857561 |
| 16 | 1058.2113 | 1062.49436 | 595.014021 | 958.968896 | 215.337117 | 990.067165 | 1099.01115 |
| 19 | 2283.67238 | 1942.13559 | 1214.42334 | 1898.93861 | 392.063742 | * | * |
| 22 | * | * | * | * | * |  |  |
| Day | PKR MS2-cGAMP (1 µg cGAMP) |  |  |  |  |  |  |
| 0 | 73.6322297 | 41.2824124 | 45.3864027 | 56.837311 | 22.101185 | 31.1127126 |  |
| 2 | 71.0906844 | 33.004366 | 133.961323 | 63.2464805 | 31.0055089 | 22.7637605 |  |
| 4 | 40.7561957 | 18.1521224 | 111.879922 | 23.3635596 | 19.0383641 | 10.3474658 |  |
| 7 | 35.9225412 | 17.4061983 | 170.714516 | 49.0542807 | 13.6887674 | 16.9603278 |  |
| 10 | 14.4802917 | 17.3431622 | 250.551313 | 66.1670375 | 10.8745842 | 7.99184519 |  |
| 13 | 9.04751667 | 6.77676407 | 430.779848 | 50.0620329 | 8.24887983 | 3.36477663 |  |
| 16 | 0 | 0 | 615.316611 | 85.4224175 | 4.16562096 | 6.44221273 |  |
| 19 | 0 | 0 | 1372.12975 | 214.581263 | 0 | 8.85234051 |  |
| 22 | 0 | 0 | 2658.02037 | 348.256855 | 0 | 10.412813 |  |
| 25 | 0 | 0 | * | 396.928218 | 0 | 15.2585773 |  |
| 27 | 0 | 0 |  | 554.695689 | 0 | 14.1494569 |  |
| 30 | 0 | 0 |  | 808.48914 | 0 | 22.5171611 |  |
| 32 | 0 | 0 |  | 1431.42378 | 0 | 34.8132448 |  |
| 33 | 0 | 0 |  | 1713.03776 | 0 | 61.0117415 |  |
| 34 | 0 | 0 |  | 2537.56283 | 0 | 100.878882 |  |
| 35 | 0 | 0 |  | * | 0 | 45.0227926 |  |
| 37 | 0 | 0 |  |  | 0 | 56.2079024 |  |
| 43 | 0 | 0 |  |  | 0 | 154.386879 |  |
| 47 | 0 | 0 |  |  | 0 | 259.665911 |  |
| 55 | 0 | 0 |  |  | 0 | 1031.37525 |  |
| 69 | 0 | 0 |  |  | 0 | 2000 |  |
| Day | Free cGAMP (50 µg) |  |  |  |  |  |  |
| 0 | 62.2423269 | 42.8009326 | 61.7776864 | 54.1679647 | 25.8244152 | 28.3569264 |  |
| 2 | 66.8365617 | 29.1437696 | 57.0599054 | 38.6877082 | 19.4034527 | 48.0887043 |  |
| 4 | 41.2835999 | 11.1450544 | 29.6641745 | 37.8296356 | 18.4805319 | 42.9275284 |  |
| 7 | 49.8600913 | 10.8999568 | 52.4029734 | 41.36148 | 22.7821597 | 79.3394899 |  |
| 10 | 64.9064163 | 8.98219039 | 64.6773267 | 61.9103967 | 26.8467942 | 109.613429 |  |
| 13 | 86.643696 | 12.8519807 | 102.177276 | 115.279046 | 83.7220829 | 167.63507 |  |
| 16 | 168.264197 | 10.448156 | 207.05546 | 195.653435 | 161.012961 | 408.442233 |  |
| 19 | 367.814326 | 18.6103299 | 428.532148 | 278.380452 | 317.958465 | 570.963931 |  |
| 22 | 546.755794 | 33.309754 | 647.117224 | 379.306409 | 493.547523 | * |  |
| 25 | 632.045025 | 34.2623289 | 969.21124 | 257.025607 | 853.224678 |  |  |
| 27 | 463.957513 | 29.568335 | 1210.93794 | 230.146221 | 1148.7698 |  |  |
| 30 | 669.934329 | 65.9936776 | 2400.06829 | 278.86762 | 848.860241 |  |  |
| 32 | 893.146283 | 178.716209 | * | 564.092291 | 1006.91588 |  |  |
| 33 | 1258.80515 | 193.851095 |  | 596.671436 | 1570.62992 |  |  |
| 34 | 1779.52332 | 245.312079 |  | 907.008984 | 1312.09305 |  |  |
| 35 | * | 225.552806 |  | 871.080005 | 1653.20242 |  |  |
| 37 |  | 353.878933 |  | 935.618137 | 2147.22024 |  |  |
| 43 |  | 1449.58353 |  | * | * |  |  |
| 47 |  | 2998.12505 |  |  |  |  |  |
| 55 |  | * |  |  |  |  |  |

| Day | ADU-S100 (50 µg) |  |  |  |  |  |
| --- | --- | --- | --- | --- | --- | --- |
| 0 | 57.3529155 | 41.8369082 | 66.700737 | 25.5413996 | 44.7218532 | 29.6368112 |
| 2 | 49.0513747 | 16.9261724 | 54.4067927 | 13.7754204 | 21.0636766 | 22.4607067 |
| 4 | 38.275858 | 24.2863333 | 19.2330124 | 25.5851557 | 16.1076932 | 15.1948721 |
| 7 | 43.7131674 | 21.4900687 | 18.2220416 | 16.9658303 | 21.9432341 | 20.2526792 |
| 10 | 62.4915322 | 13.9626201 | 15.7675614 | 15.1281185 | 7.6785551 | 17.5961359 |
| 13 | 69.2483346 | 0 | 9.23320783 | 12.070355 | 10.088075 | 11.9102888 |
| 16 | 115.750836 | 0 | 5.20991254 | 9.86616126 | 9.51836339 | 4.82792733 |
| 19 | 183.659014 | 0 | 13.9571537 | 8.24219871 | 6.74647807 | 3.14391063 |
| 22 | 373.754107 | 0 | 31.07578 | 11.3352888 | 4.49993354 | 2.06793033 |
| 25 | 391.522367 | 0 | 35.4336214 | 0 | 0 | 0 |
| 27 | 308.277199 | 0 | 60.6463539 | 0 | 0 | 0 |
| 30 | 339.115787 | 0 | 138.418058 | 0 | 0 | 0 |
| 32 | 652.6298 | 0 | 214.564998 | 0 | 0 | 0 |
| 33 | 810.453815 | 0 | 258.913228 | 0 | 0 | 0 |
| 34 | 1234.03395 | 0 | 210.443266 | 0 | 0 | 0 |
| 35 | 1262.80391 | 0 | 142.490881 | 0 | 0 | 0 |
| 37 | 1448.6075 | 0 | 210.589584 | 0 | 0 | 0 |
| 43 | * | 0 | 1125.3067 | 0 | 0 | 0 |
| 47 |  | 0 | 272.240215 | 0 | 0 | 0 |
| 55 |  | 0 | 1616.40892 | 0 | 0 | 0 |
| 69 |  | 0 | * | 0 | 0 | 0 |

| Day | PKR MS2-cGAMP (5 µg cGAMP) |
| --- | --- |

|  |  |  |  |  |  |  |
| --- | --- | --- | --- | --- | --- | --- |
| 0 | 41.6242303 | 68.914033 | 29.924047 | 45.2144717 | 25.2072912 | 57.1914533 |
| 2 | 45.3826265 | 35.2180076 | 32.9980096 | 61.7274702 | 26.7094186 | 26.780227 |
| 4 | 28.7458555 | 26.7094228 | 28.2445956 | 33.5091153 | 17.9095914 | 9.21329432 |
| 7 | 41.2949285 | 24.7440006 | 35.3860781 | 26.3879792 | 31.4778185 | 33.5105991 |
| 10 | 33.350978 | 59.3968147 | 35.6760644 | 23.3602782 | 25.9968363 | 34.5169424 |
| 13 | 24.0748099 | 103.617278 | 30.0948868 | 4.90728292 | 3.96113993 | 17.6416596 |
| 16 | 18.0802977 | 268.966479 | 45.0449487 | 0 | 2.29679745 | 8.54051231 |
| 19 | 44.0222477 | 579.530099 | 168.596096 | 0 | 0 | 0 |
| 22 | 62.8387017 | 704.522208 | 177.34102 | 2.56856615 | 0 | 0 |
| 25 | 89.9645029 | 2030.61997 | 365.631848 | 0 | 0 | 0 |
| 27 | 180.006624 | * | 401.109569 | 0 | 0 | 0 |
| 30 | 295.872264 |  | 749.922357 | 0 | 0 | 0 |
| 32 | 443.506753 |  | 1117.90872 | 0 | 0 | 0 |
| 33 | 553.223722 |  | 1429.55269 | 0 | 0 | 0 |
| 34 | 815.176105 |  | 2762.86958 | 0 | 0 | 0 |
| 35 | 832.725503 |  | * | 0 | 0 | 0 |
| 37 | 1048.02372 |  |  | 0 | 0 | 0 |
| 43 | * |  |  | 0 | 0 | 0 |
| 47 |  |  |  | 0 | 0 | 0 |
| 55 |  |  |  | 0 | 0 | 0 |
| 69 |  |  |  | 0 | 0 | 0 |
| Day | **PKR MS2-cGAMP (0.2 µg cGAMP)** |  |  |  |  |  |
| 0 | 42.2225361 | 31.7740943 | 16.1766627 | 46.3015491 | 103.886971 | 39.0693908 |
| 2 | 69.9221449 | 49.1252508 | 14.5635858 | 41.7331556 | 85.0620318 | 45.652206 |
| 4 | 35.3298797 | 49.3230225 | 11.0871077 | 20.6171343 | 76.6463491 | 49.309433 |
| 7 | 49.5222994 | 86.7529888 | 23.2355648 | 38.9884555 | 89.187687 | 147.335401 |
| 10 | 48.3095059 | 112.05375 | 30.1016727 | 43.0452983 | 131.192514 | 372.053741 |
| 13 | 81.267278 | 198.01292 | 65.1535948 | 98.6896398 | 230.249403 | 678.059996 |
| 16 | 231.49543 | 294.366996 | 220.221247 | 137.033651 | 382.784432 | * |
| 19 | 419.582177 | 491.685489 | * | 322.60743 | 581.551027 |  |
| 22 | 653.337 | 739.773827 |  | 509.028601 | 789.178655 |  |
| 25 | 722.290671 | 783.844165 |  | 921.49081 | 1010.30469 |  |
| 27 | 947.457365 | 1313.30511 |  | 924.870005 | 1143.27743 |  |
| 30 | * | 823.180062 |  | 1415.00299 | 2221.28018 |  |
| 32 |  | 1533.24364 |  | * | 2010.18265 |  |
| 33 |  | * |  |  | * |  |

**Table S2.** Tumor volumes in mice from the *in vivo* tumor response study represented in **Figure 4e**. All values are given in mm^3^. Each column represents the tumor volumes of one mouse in each sample. Mice that were sacrificed or died during the study are marked with an asterisk (*), while mice with no measurable tumor volume are indicated with a zero (0). The study was concluded 59 days after STING agonist or control treatment.

| Day | PBS – STING WT mice |  |  |  |  |  |  |
| --- | --- | --- | --- | --- | --- | --- | --- |
| 0 | 10.5623157 | 7.2763815 | 75.7625856 | 20.0544195 | 41.7534942 | 214.652774 | 74.2871518 |
| 2 | 28.2256612 | 24.641082 | 189.401951 | 23.1873058 | 81.781442 | 403.045306 | 162.527871 |
| 4 | 19.6768943 | 41.0051056 | 334.095863 | 19.5595433 | 112.040211 | 558.626523 | 226.828714 |
| 7 | 24.6490978 | 67.7962826 | 414.994336 | 54.4096202 | 150.710041 | 1067.38715 | 269.74337 |
| 12 | 148.071636 | 228.791141 | 1683.64885 | 123.517412 | 361.734288 | 1560.77209 | 851.034911 |
| 16 | 325.999206 | 484.323332 | * | 275.277669 | 670.321366 | * | * |
| 19 | 665.780957 | 765.440561 |  | 660.056159 | 1277.79921 |  |  |
| 22 | 1342.94715 | 1678.4412 |  | 1438.10013 | 2096.0586 |  |  |
| 26 | 2656.62328 | 2777.87968 |  | 2083.73775 | * |  |  |
| 29 | * | * |  | * |  |  |  |
| Day | PKR MS2-cGAMP (2 µg) – STING WT mice |  |  |  |  |  |  |
| 0 | 40.8176326 | 140.4307 | 63.2619329 | 27.2285605 | 12.6868161 |  |  |
| 2 | 51.0185976 | 120.965526 | 73.8720893 | 22.5182481 | 22.8280689 |  |  |
| 4 | 68.1537194 | 148.023544 | 105.724699 | 19.665038 | 9.14206918 |  |  |
| 7 | 166.893905 | 149.499114 | 65.8069078 | 5.93984274 | 7.31020964 |  |  |
| 12 | 401.174062 | 267.564772 | 78.6283778 | 17.1148077 | 1.49527244 |  |  |
| 16 | 1099.98338 | 511.521153 | 97.5735481 | 85.1725572 | 0 |  |  |
| 19 | * | 557.802222 | 111.778501 | 111.556896 | 0 |  |  |
| 22 |  | 2754.87832 | 333.299628 | 308.112194 | 0 |  |  |
| 26 |  | * | 305.940922 | 518.447179 | 0 |  |  |
| 29 |  |  | 616.60313 | 1030.01174 | 0 |  |  |
| 33 |  |  | 1274.28896 | 1968.92383 | 0 |  |  |
| 36 |  |  | 1364.10275 | 2907.97493 | 0 |  |  |
| 41 |  |  | 3500.15493 | * | 0 |  |  |
| 45 |  |  | * |  | 0 |  |  |
| 49 |  |  |  |  | 0 |  |  |
| 53 |  |  |  |  | 0 |  |  |
| 55 |  |  |  |  | 0 |  |  |
| 59 |  |  |  |  | 0 |  |  |
| Day | PBS – STING<sup>gt/gt</sup> mice |  |  |  |  |  |  |
| 0 | 29.9768258 | 56.0446935 | 27.1403655 | 45.45761 | 96.5662069 |  |  |
| 2 | 81.6102519 | 123.91535 | 75.4917049 | 89.3109181 | 156.24945 |  |  |
| 4 | 118.480392 | 227.297496 | 122.601439 | 114.843345 | 250.879212 |  |  |
| 7 | 223.93459 | 350.614876 | 227.2191 | 250.068783 | 360.25177 |  |  |
| 12 | 437.115069 | 1012.97789 | 724.54677 | 526.020302 | 1104.07449 |  |  |
| 16 | 561.101426 | * | 1259.51752 | 1386.19644 | * |  |  |
| 19 | * |  | * | * |  |  |  |
|  | **PKR MS2-cGAMP (2 µg) – STING<sup>gt/gt</sup> mice** |  |  |  |  |  |  |
| 0 | 34.7887655 | 91.305476 | 14.4942707 | 38.1326516 | 77.9818506 |  |  |
| 2 | 80.4151796 | 147.57918 | 46.3261431 | 29.2910789 | 168.198514 |  |  |
| 4 | 152.519071 | 204.410918 | 77.3570381 | 35.7949606 | 211.889668 |  |  |
| 7 | 272.760746 | 271.684933 | 106.903108 | 22.6298878 | 217.742782 |  |  |
| 12 | 720.530659 | 974.818003 | 406.303544 | 158.379555 | 599.588201 |  |  |
| 16 | 1101.26263 | 2396.43153 | 531.960116 | 240.85363 | 1080.5039 |  |  |
| 19 | 2191.9424 | * | 1084.47352 | 408.515845 | 1755.665 |  |  |
| 22 | * |  | 921.758817 | 327.829152 | 1802.4092 |  |  |
| 26 |  |  | 1912.35986 | 1052.78442 | 3623.40168 |  |  |
| 29 |  |  | 2395.85191 | 1596.44215 | * |  |  |
| 33 |  |  | * | 2134.94474 |  |  |  |
| 36 |  |  |  | * |  |  |  |

## MATERIALS AND METHODS

All reagents were obtained from commercial sources and used without further purification unless otherwise indicated. Dulbecco’s phosphate-buffered saline (DPBS) and all mammalian cell growth media was purchased from Gibco. All other aqueous buffers and media were prepared using milli-Q H_2_O purified to a resistivity of 18.2 MΩ at 25 °C obtained from a MilliporeSigma Milli-Q EQ7000 purification system. Free 2’,3’-cGAMP was generated in-house or obtained from APExBIO. ADU-S100 was obtained as a gift from Novartis. STF-1084 and endocytosis inhibitors were obtained from Sigma-Aldrich. No unexpected or unusually high safety hazards were encountered.

### Equipment and Instrumentation

Protein purification was performed using a Cytiva AKTA go fast protein liquid chromatograph (FPLC). Colorimetric 96-well plate measurements were performed on an Agilent BioTek Synergy H1 plate reader. Small molecules were purified on a Biotage Isolera One flash chromatography system. Dynamic light scattering (DLS) was performed on a Horiba nanoPartica SZ-100V2. Flow cytometry measurements were performed on a ThermoFisher Attune NxT flow cytometer at the QB3 Cell and Tissue Analysis Facility (CTAF), UC Berkeley. RT-qPCR was performed using a BioRad CFX Connect thermocycler, also part of QB3 CTAF, UC Berkeley.

### Liquid Chromatography/Mass Spectrometry (LC/MS)

Purified proteins and small molecules were analyzed by electrospray ionization time-of-flight mass spectrometry (ESI-TOF-MS). Samples were first separated with an elution gradient of milli-Q H_2_O + 0.1% (v/v) formic acid and Optima MS grade MeCN + 0.1% (v/v) formic acid, on an Agilent PLRP-S 1000 Å monolithic analytical column using an Agilent 1260 series liquid chromatography (LC) system, then mass spectra were obtained on an Agilent 6530 Q-TOF MS system.

### Mammalian Cell Culture

THP-1 reporter cells were cultured in RPMI + GlutaMAX supplemented with 10% FBS. MC38 cells were cultured in DMEM + GlutaMAX supplemented with 10% FBS. All cells were maintained at 37 °C in a humidified environment and 5% CO_2_.

### *In vivo* Mouse Experiments

All studies were performed according to protocols approved by the UC Berkeley Animal Care and Use Committee (ACUC). C57BL/6J and STING^gt/gt^ mice were purchased from Jackson Laboratory and maintained at UC Berkeley.

### Protein Preparation and Sequences

#### Expression and Purification of MS2 Constructs

MS2 capsid sequences were expressed and purified based on a protocol adapted from a previous work^39,43^. Briefly, a single colony of DH10b *E. coli* containing a pBAD plasmid containing the gene for each MS2 variant was grown overnight in a 10 mL culture of LB at 37 °C, then subcultured into 1 L of 2xYT media with 20 µg/mL chloramphenicol and incubated shaking at 37 °C. OD_600_ was monitored until reaching a value of 0.6, upon which 0.1% (w/v) arabinose was added and cells grown overnight. The next day, cells were collected by centrifugation and resuspended in 20 mL of 10 mM sodium phosphate, pH 7.4 + 0.02% (w/v) NaN_3_, and lysed on ice by sonication at 75% amplitude for 10 min (2 s on, 4 s off). The cell lysate was then centrifuged at 14,000 x g for 30 min and supernatant collected. MS2 protein was precipitated by addition of an equal volume of saturated aqueous (NH_4_)_2_SO_4_ and rotation overnight at 4 °C. The mixture was then centrifuged at 14,000 x g for 30 min and supernatant was discarded. The precipitate was redissolved in 10 mM sodium phosphate, pH 7.4 + 0.02% NaN_3_, centrifuged again to remove undissolved material, and filtered through a 0.22 µm membrane filter. Sample was then purified by FPLC first using two HiScreen CaptoCore 700 columns connected in series, with an isocratic flow of the same phosphate buffer. The flow-through was then desalted into 20 mM sodium phosphate buffer, pH 7.4, using a HiPrep 26/10 Desalting column. For PKR MS2, samples were further purified by FPLC on a 5 mL HiTrap Heparin HP affinity column, eluting with a gradient of 20 mM sodium phosphate, pH 7.4 (buffer A) to 20 mM sodium phosphate + 2 M NaCl, pH 7.4 (buffer B). The purified MS2 was then buffer exchanged into phosphate-buffered saline (PBS) using an Amicon 100kDa MWCO spin concentrator. Purified MS2 was confirmed by SDS-PAGE and LC/MS and stored at 4 °C.

#### Expression and Purification of SUMO-cGAS

A pET-29b(+) plasmid vector containing the gene for His6-SUMO-cGAS was purchased from Twist Biosciences and transformed into BL21(DE3)* *E. coli* cells. The transformed cells were plated on kanamycin-containing agar overnight, and a single colony was inoculated into 5 mL LB media and grown overnight at 37 °C. The sample was then subcultured into 1 L of TB media with 50 µg/mL kanamycin and grown at 37 °C until OD_600_ reached 0.6, at which point the sample was cooled to 16 °C. Once equilibrated at 16 °C, 0.1 mM IPTG was added and cells were grown for 16 h. Cells were then harvested by centrifugation and dissolved in freshly prepared lysis buffer (50 mM Tris, 300 mM NaCl, 30 mM imidazole, 1 mM TCEP, pH 8.0) with PMSF added to 1 mM. Cells were sonicated on ice at 75% amplitude for 10 min (2 s on, 4 s off), then centrifuged at 16,000 x g for 30 min. The supernatant was collected and filtered through a 0.22 µm PES syringe filter. An AKTA go FPLC was first cleaned by flushing all lines with 70% acetic acid in water, then 2 N NaOH, then milli-Q H_2_O to remove all potential contaminants. The sample was then purified on a 5 mL HisTrap HP, using a gradient of lysis buffer (buffer A) and the same buffer with 300 mM imidazole (buffer B). The purified protein was then desalted into 50 mM Tris, 1 mM TCEP, pH 8.0 without added NaCl using a HiPrep 26/10 desalting column. Care was taken to minimize NaCl concentration from pH adjustment, as this was observed to inhibit cGAS activity. Purified SUMO-cGAS was confirmed by LC/MS. Protein was either used immediately or snap-frozen in 10% glycerol and stored at −80 °C.

#### Protein Sequences

For MS2 sequences, engineered residues are underlined.

**MS2 N87C –** molecular weight: 13,717 Da

ASNFTQFVLVDNGGTGDVTVAPSNFANGVAEWISSNSRSQAYKVTCSVRQSSAQNRKYT IKVEVPKVATQTVGGVELPVAAWRSYL<u>C</u>MELTIPIFATNSDCELIVKAMQGLLKDGNPIPS AIAANSGIY

**MS2 S37P T71K G73R N87C (PKR MS2)** – molecular weight: 13,854 Da

ASNFTQFVLVDNGGTGDVTVAPSNFANGVAEWISSN<u>P</u>RSQAYKVTCSVRQSSAQNRKYT IKVEVPKVATQ<u>K</u>V<u>R</u>GVELPVAAWRSYL<u>C</u>MELTIPIFATNSDCELIVKAMQGLLKDGNPIPS AIAANSGIY

**SUMO-cGAS** – molecular weight: 55,740 Da (truncated human cGAS sequence <u>underlined</u>)

GSSHHHHHHSSGLVPRGSHMSDSEVNQEAKPEVKPEVKPETHINLKVSDGSSEIFFKIKKTTPLRRLMEAFAKRQGKEMDSLRFLYDGIRIQADQTPEDLDMEDNDIIEAHREQIGGMGASKLRAVLEKLKLSRDDISTAAGMVKGVVDHLLLRLKCDSAFRGVGLLNTGSYYEHVKISAPNEFDVMFKLEVPRIQLEEYSNTRAYYFVKFKRNPKENPLSQFLEGEILSASKMLSKFRKIIKEEINDIKDTDVIMKRKRGGSPAVTLLISEKISVDITLALESKSSWPASTQEGLRIQNWLSAKVRKQLRLKPFYLVPKHAKEGNGFQEETWRLSFSHIEKEILNNHGKSKTCCENKEEKCCRKDCLKLMKYLLEQLKERFKDKKHLDKFSSYHVKTAFFHVCTQNPQDSQWDRKDLGLCFDNCVTYFLQCLRTEKLENYFIPEFNLFSSNLIDKRSKEFLTKQIEYERNNEFPVFD EF

### Experimental Protocols

#### Enzymatic Synthesis and Purification of 2’,3’-cGAMP

In a 500 mL solution of 50 mM Tris, pH 8.0, without added NaCl was added 2 mM ATP, 2 mM GTP, 10 mM MgCl_2_, 0.1 mg/mL herring testis DNA, 1 mM TCEP, and 1 µM cGAS. The sample was stirred for 24 h at 37 °C, after which the solution usually turned uniformly cloudy. The sample was then frozen and lyophilized and redissolved in 10 mL milli-Q water. The cloudy solution was centrifuged to remove precipitate, and the supernatant was collected, filtered through a 0.22 µm PES syringe filter, then purified on a 15.5 g ISCO C18aq flash chromatography cartridge, using a gradient of 50 mM triethylammonium acetate (TEAA) in water and MeCN. Purified cGAMP eluted at approximately 15% MeCN, and fractions were collected, verified using LC/MS, and lyophilized to obtain pure cGAMP.

#### Construction of MS2-cGAMP

MS2 N87C was diluted in DPBS to 50 µM, and 3 equiv. cGAMP-disulfide was added. The sample was incubated at 4 °C for at least 1 h, after which complete conjugation was confirmed by LC/MS. Excess cGAMP was then removed by buffer exchanging into clean DPBS sequentially 5 times using a 0.5 mL Amicon 100 kDa MWCO spin concentrator. The resulting samples were then filtered through a 0.22 µm cellulose acetate centrifuge filter.

#### THP-1 STING reporter cell experiments

THP-1 cells bearing a tdTomato STING reporter and, where applicable, a SLC19A1 knockout were generated in a previous study^19^. For STING KD reporter cells, THP-1 monocytes were transduced with 1) a lentiviral dCas9-HA-BFP-KRAB-NLS expression vector (Addgene, plasmid no.102244), 2) a lentiviral vector (pCRISPRia-v2, Addgene, plasmid no. 84832) expressing a control gRNA (GGAGAGACGGTACCGTCTCA) or STING-targeting gRNA (GGCTGCTCTGGATGATGACG) and 3) a lentiviral vector encoding the tdTomato reporter gene driven by the ISREs and the minimal mouse IFN-β promoter.

For STING agonist treatments, THP-1 STING reporter cells were spun down, resuspended in fresh media with or without FBS or inhibitor, and plated in a 48-well plate at 50,000 cells/well and treated with a 10x dilution of MS2-cGAMP, free cGAMP, or free ADU-S100 in DPBS for 24 h. Cells were then transferred to a V-bottom 96-well plate and analyzed by flow cytometry. Cells were gated first by FSC/SSC, then doublets removed by FSC-A/FSC-H gating, followed by presence of a constitutive GFP signal in cells with the STING reporter, followed by tdTomato analysis.

#### THP-1 STING reporter cell uptake of fluorescently labeled MS2

AZdye 647-maleimide was obtained from Vector Labs and dissolved to 10 mM in DMSO. PKR MS2 was incubated with 3 equiv of AZdye 647-maleimide at pH 7.2 for 1 h at 4 °C, then analyzed by LC/MS to confirm complete modification. Samples were then desalted using a 0.5 mL Zeba 40 kDa spin desalting column into DPBS. THP-1 cells were then plated at 50,000 cells/well in a 48-well plate and treated with 0.2 µM MS2-dye for 24 h. Cells were then transferred to a V-bottom 96-well plate and analyzed by flow cytometry using identical gating as before and analysis of the AlexaFluor 647 channel.

#### THP-1 STING reporter endocytosis inhibitor screen

THP-1 cells were plated in a 48-well plate at 50,000 cells/well and pre-incubated with 10 U/mL heparin, 1 U/mL heparinase, 50 µM chloroquine, 5 µM cytochalasin D, or 80 µM dynasore for 2 h. Cells treated with heparinase were spun down, media was removed by aspiration, and fresh media was replaced. Cells were then treated with fluorescently labeled MS2 for uptake or STING agonists for STING activation and analyzed as described above.

#### RT-qPCR of STING agonist-treated cells

RT-qPCR conditions were adopted from a previous study^40^. Briefly, THP-1 or MC38 cells were treated with 100 nM PKR-cGAMP, 100 µM free cGAMP, or 10 µM free ADU-S100 for 4 or 24 h. Adherent MC38 cells were then lifted by 0.25% trypsin. All cell types were then centrifuged, and media was removed by aspiration. RNA was extracted using TriZOL buffer and chloroform per manufacturer instructions, and the resulting RNA was converted to cDNA using a BioRad iScript cDNA synthesis kit. A 350 ng sample of cDNA was mixed with SYBR Green Master Mix and 500 nM of primer in 20 µL nuclease-free water. qPCR data was obtained using the following conditions on a thermocycler: 95 °C for 2 min, followed by 42 repeats of 95 °C for 15 s, followed by 60 °C for 60 s. Primer sequences: Mouse *IFNB* fwd: 5’-ATAAGCAGCTCCAGCTCCAA-3’, rev: 5’-CTGTCTGCTGGTGGAGTTCA-3’; Human *IFNB* fwd: 5’-AAACTCATGAGCAGTCTGCA-3’, rev: 5’-AGGAGATCTTCAGTTTCGGAGG-3’; *ACTB* (endogenous housekeeping control) fwd: 5’-AGAGCTACGAGCTGCCTGAC-3’, rev: 5’-AGCACTGTGTTGGCGTACAG-3’.

#### Cell viability assay

THP-1 cells were treated with STING agonists or controls as described above. After 24 h, samples were split in half, with half the samples analyzed by flow cytometry to ensure expected STING activation. The other half of each well was transferred to a 96-well U-bottom plate, where cells were centrifuged and media removed by aspiration. MTS reagent was obtained from APExBIO (K2250) and diluted 10x in DPBS. 200 µL diluted reagent was added to each well and cells were incubated and analyzed according to the manufacturer’s instructions.

#### PMA differentiation of THP-1 STING reporter cells

THP-1 cells were differentiated based on a previous study^40^. Cells were mixed with 125 ng/mL of phorobol 12-myristate 13-acetate and plated in a 48-well plate at 50,000 cells/well. Cells were incubated for 48 h, after which differentiated cells had adhered to the plate bottom. Media was aspirated and replaced with PMA-free RPMI media, in which cells were incubated for 24 h. Cells were then treated with STING agonists and controls as previously. Cells were lifted with 40 µL 0.25% trypsin and diluted with RPMI, after which they were analyzed by flow cytometry.

#### *In vivo* antitumor studies

For *in vivo* tumor inoculation, cells were washed and resuspended in PBS, and 50 μL containing 4 x 10^6^ cells was injected subcutaneously into C57BL/6J and STING^gt/gt^mice. Tumor dimensions (length, width, and height) were measured using digital calipers, and tumor volume was calculated using the ellipsoid formula: V = (π/6) × length × width × height. When tumors reached a volume of 50 mm^3^, they were injected intratumorally with PBS, PKR MS2-cGAMP corresponding to 0.2, 1, 2, or 5 μg cGAMP, 50 μg ADU-S100, or 50 μg free cGAMP in 50 μL PBS. In all experiments, prior to treatment initiation, tumor-bearing mice were stratified by tumor volume and randomly assigned to treatment groups such that mean tumor sizes were comparable across groups. Mice were euthanized upon reaching institutional humane endpoints.

## Synthetic Methods

### Synthesis of cGAMP-disulfide

**Scheme S1.**
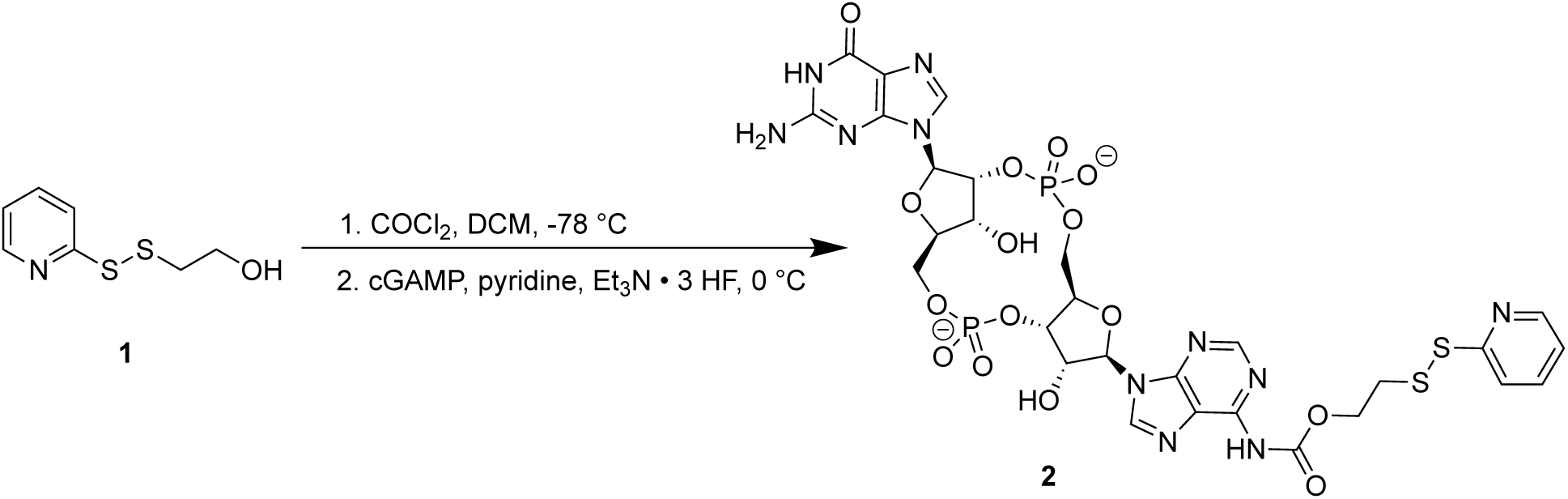
Synthesis of cGAMP-disulfide conjugate.

#### cGAMP-disulfide conjugate 2

Synthesis of the pyridyl disulfide compound **1** was performed as in a previous study^40^. Compound **1** (42 mg, 0.223 mmol) was dissolved in 1 mL of anhydrous DCM. A solution of phosgene (15% in toluene, 0.54 mL, 0.742 mmol) was added dropwise and the reaction was stirred at 0 °C under N_2_ for 1 h. Solvent and excess phosgene were then removed by vacuum. About 1 mL of dry DCM was added and then removed under vacuum 3 times to azeotropically remove any excess phosgene. A portion of 2’,3’-cGAMP (5 mg, 0.00742 mmol) was suspended in 1 mL of dry pyridine, and triethylammonium trihydrofluoride (Et_3_N • 3 HF) was added in 25 µL increments until the solids were fully dissolved, with roughly 100-150 µL HF-TEA needed. The solution was then added to the reaction mixture, and the resulting reaction was stirred for 90 min at 0 °C under N_2_. The volatile components were then evaporated under vacuum, and the reaction was quenched with 500 µL water, then dissolved in 1.5 mL DMSO. The product was purified twice by Biotage C18 column chromatography (50 mM NEt_3_•AcOH in H_2_O/MeCN) with elution at 30% MeCN. Concentration of the fractions yielded **2** as a white solid (0.7 mg, 11% yield).

